# Tumour gene expression signature in primary melanoma predicts long-term outcomes: A prospective multicentre study

**DOI:** 10.1101/2020.02.24.961771

**Authors:** Manik Garg, Dominique-Laurent Couturier, Jérémie Nsengimana, Nuno A. Fonseca, Matthew Wongchenko, Yibing Yan, Martin Lauss, Göran B Jönsson, Julia Newton-Bishop, Christine Parkinson, Mark R. Middleton, Tim Bishop, Pippa Corrie, David J. Adams, Alvis Brazma, Roy Rabbie

**Affiliations:** European Molecular Biology Laboratory, European Bioinformatics Institute (EMBL-EBI), Hinxton, Cambridgeshire, UK; Cancer Research UK Cambridge Institute, University of Cambridge, Li Ka Shing Centre, Robinson Way, Cambridge, UK; University of Leeds School of Medicine, Leeds, United Kingdom; CIBIO/InBIO-Centro de Investigação em Biodiversidade e Recursos Genéticos, Universidade do Porto, Rua Padre Armando Quintas, 4485-601 Vairão, Portugal; Oncology Biomarker Development, Genentech, Inc. 1 DNA Way, South San Francisco, CA 94080, USA; Lund University Cancer Center, Lund University, Lund, Sweden; Cambridge Cancer Centre, Cambridge University Hospitals NHS Foundation Trust, Cambridge, UK; Oxford NIHR Biomedical Research Centre and Department of Oncology, University of Oxford, Oxford, UK; Experimental Cancer Genetics, The Wellcome Sanger Institute, Hinxton, Cambridgeshire, UK

**Keywords:** Melanoma, prognostic biomarkers, adjuvant therapy, immune response, precision medicine

## Abstract

**Purpose:** Predicting outcomes after resection of primary melanoma remains crude, primarily based on tumour thickness. We explored gene expression signatures for their ability to better predict outcomes.

**Methods:** Differential expression analysis of 194 primary melanomas resected from patients who either developed distant metastasis (n=89) or did not (n=105) was performed. We identified 121 metastasis-associated genes that were included in our prognostic signature, “Cam_121”. Several machine learning classification models were trained using nested leave-one-out cross validation (LOOCV) to test the signature’s capacity to predict metastases, as well as regression models to predict survival. The prognostic accuracy was externally validated in two independent datasets.

**Results:** Cam_121 performed significantly better in predicting distant metastases than any of the models trained with the clinical covariates alone (p_Accuracy_=4.92×10^−3^), as well as those trained with two published prognostic signatures. Cam_121 expression score was strongly associated with progression-free survival (HR=1.7, p=3.44×10^−6^), overall survival (HR=1.73, p=7.71×10^−6^) and melanoma-specific survival (HR=1.59, p=0.02). Cam_121 expression score also negatively correlated with measures of immune cell infiltration (ρ=−0.73, p<2.2×10^−16^), with a higher score representing reduced tumour lymphocytic infiltration and a higher absolute 5-year risk of death in stage II melanoma.

**Conclusions:** The Cam_121 primary melanoma gene expression signature outperformed currently available alternatives in predicting the risk of distant recurrence. The signature confirmed (using unbiased approaches) the central prognostic importance of immune cell infiltration in long-term patient outcomes and could be used to identify stage II melanoma patients at highest risk of metastases and poor survival who might benefit most from adjuvant therapies.

**Translational relevance:** Predicting outcomes after resection of primary melanoma is currently based on traditional histopathological staging, however survival outcomes within these disease stages varies markedly. Since adjuvant systemic therapies are now being used routinely, accurate prognostic information is needed to better risk stratify patients and avoid unnecessary use of high cost, potentially harmful drugs, as well as to inform future adjuvant strategies. The Cam_121 gene expression signature appears to have this capability and warrants evaluation in prospective clinical trials.

## Background

Melanoma is among the few cancers demonstrating an increasing incidence over time (1). Most patients present with primary tumours and the majority will be cured by local surgery. Outcomes from metastatic melanoma have improved radically in the last 10 years with the introduction of new systemic therapies (2), although median survival remains around 3 years. It is the large number of patients presenting with early stage disease who unfortunately experience disease recurrence over time and represent the majority of deaths due to melanoma (3). Optimal management of early melanoma is therefore key to improving outcomes.

Patients with resected AJCC stage III melanoma are now eligible for adjuvant immune checkpoint inhibitors as well as *BRAF* targeted therapies, based on randomised trials confirming reduction in risk of relapse and improved overall survival (4-7). Clinical trials are underway to evaluate similar therapies in resected stage IIB/C patients (8), whose outcomes reflect that of stage IIIA/B melanoma, untreated (9). As such the number of patients eligible for treatment with adjuvant therapies over the coming years is expected to increase substantially. These modern anticancer drugs are high cost and carry risk of both life-changing and life-threatening toxicities, so there is growing desire to more accurately predict those patients at high-risk of recurrence in whom intervention is expected to be beneficial and to avoid over-treating patients likely already to be cured of their disease by surgery alone.

Gene expression signatures have the potential to improve the prediction of the biological behaviour of melanocytic lesions by objectively defining ‘high-risk’ on a molecular level (10). Previous transcriptomic analyses of cutaneous melanoma (CM) identified patterns of gene expression associated with survival independent of AJCC stage (11). Building on these data, Gerami *et al* first reported a prognostic gene expression profile (GEP) test utilising the 31-gene panel (28 discriminating and 3 control genes) for use in patients with CM (Decision-Dx Melanoma™) (12). The test measures the expression of individual genes from formalin-fixed paraffin-embedded (FFPE) primary melanomas to provide a binary classification of low (class 1) or high (class 2) risk for developing metastases within 5 years of diagnosis (with A and B subclasses to further stratify risk) (13-15). A further recent unsupervised clustering analyses based on 677 primary melanoma transcriptomes embedded within a population-controlled cohort study from the Leeds Melanoma Cohort (LMC) identified a six-class 150-gene prognostic signature (herein reference to as LMC_150) (16). The signature uniquely demonstrated prognostic relevance in patients with stage I primary melanoma and further predicted poor outcomes in patients undergoing immunotherapy (16). However, owing at least in-part to a lack of prospective reproducible data, proving the clinical utility of such prognostic molecular tools remains a considerable challenge and there are currently no established prognostic biomarkers able to accurately identify truly high-risk patients (17).

Using patient samples and long-term clinical outcome data from one of the largest adjuvant melanoma trials (18), we undertook RNA sequencing of the primary tumour matched with robust prospective clinical data to uncover a molecular signature that could be used to predict patient outcomes. This was then validated in two externally acquired datasets.

## Results

### 1. Prognostic signature generated using covariate corrected differential expression

The structure of the datasets and analyses are depicted in (Supplementary figure S1). Principal component analysis (PCA) showed that primary cutaneous melanomas (n=204) and melanoma spread to local lymph nodes (n=177) clustered separately, suggesting an impact of the microenvironment on tumour gene expression (Supplementary figure S2; see Supplementary methods). We therefore decided to treat these as separate datasets, focussing our analyses on the primary melanoma samples and validated our results in the lymph node metastases (see Methods). We conducted a differential expression analysis, identifying differences in gene expression levels in primary tumours between those patients with and without distant metastasis over a minimum of six years follow-up while controlling for a set of variables (Supplementary figure S3(A); Supplementary table S1). Our analyses revealed 197 significantly differentially expressed genes (DEGs, FDR adjusted p-value<0.1) associated with metastases (Supplementary figure S3(B)). These DEGs were further filtered to remove pseudogenes (n=39) and those genes not identified within the Leeds Melanoma Cohort DASL array (n=37) to enable external validation of our signature (Supplementary figure S1). We were therefore left with 121 DEGs which made up our core prognostic signature herein referred to as “Cam_121” (Supplementary table S2).

### 2. Cam_121 predicts metastases significantly better than both clinical covariates and published prognostic signatures

We trained machine learning models with the prognostic signature Cam_121 employing multiple (n=8) machine learning classifiers (see Methods). The models were evaluated by their capacity to predict whether a primary melanoma would ultimately metastasise to distant body sites, or would not. We trained the same models with the key clinical covariates, of which a number were independently associated with distant metastases (Stage AJCC 7th edition (19) (herein referred to as ‘stage’), Breslow thickness, ECOG; Eastern Cooperative Oncology Group Performance Status and the experimental adjuvant therapy) (Supplementary table S1) for comparison. We found that the models trained with both Cam_121 and clinical covariates significantly out-performed the models trained with the clinical covariates alone (p_Accuracy_=2.94×10^−3^, p_kappa_=3.23×10^−3^; Supplementary table S3), and this remained consistent across all machine learning classifiers (Supplementary figure S4; Supplementary table S3). By selecting the top performing classifier for each signature, we were able to compare the performance of the signature to baseline clinical covariates across multiple measures of model efficacy. The prognostic signature added incremental benefit across all key performance metrics including: accuracy (66.5% using the clinical covariates only vs 72.7% using the Cam_121 gene expression + clinical covariates), kappa (0.32 vs 0.45), sensitivity (60.7% vs 70.8%), specificity (71.4% vs 74.3%) (Figure 1A-D). In particular, we found that adding the signature to the clinical covariates correctly predicted an additional 42 cases (Figure 1E). We then tested the performance of published prognostic signatures from Gerami *et al* (12) (Decision-Dx Melanoma™; n=27 genes) and Thakur *et al* (16) (LMC_150; n=150 genes) using the same machine learning models in our dataset and were not able to deduce that these signatures perform better than the baseline clinical covariates at the 5% significance level (Supplementary figure S4 & Supplementary table S3).

**Figure 1.**
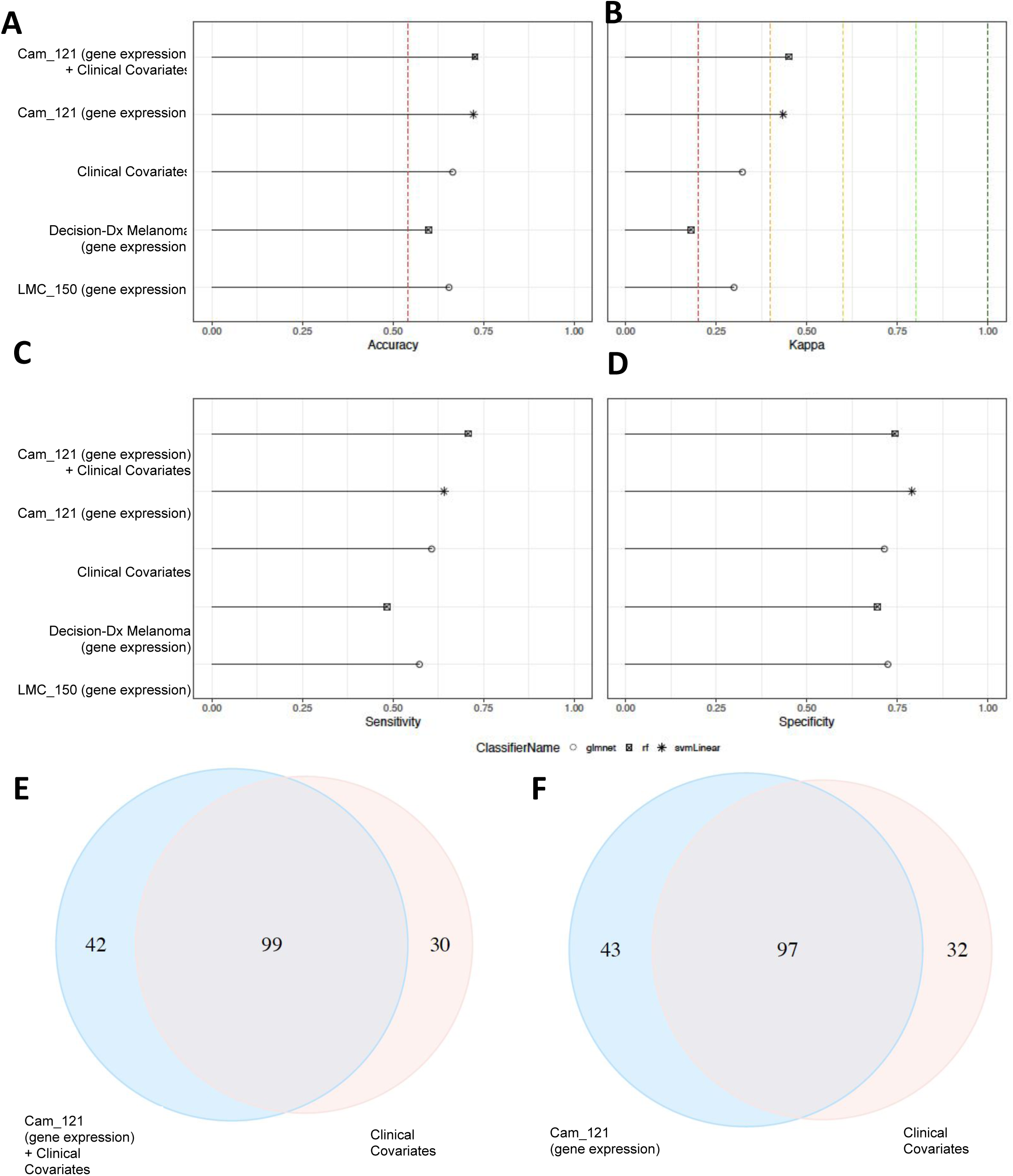
Cam_121 gene expression is strongly associated with both metastases and survival across multiple machine-learning and Cox regression analyses. Barplots indicating, for each set of predictors (y-axes), the scores of **A)** accuracy, **B)** kappa, **C)** sensitivity and **D)** specificity (x-axes) of the top performing classifier (among the 8 considered). The dashed red line in (A) represents the accuracy achieved when the classifier only predicts the majority class (metastases=“No” in our case, 105/194 (54.12%)). Coloured lines in (B) represent the range of agreement bands of [cite whoever] defined for Cohen’s Kappa values. Venn diagrams comparing the number of correctly predicted relapse outcomes (yes/no) of 194 patients of **E)** “Cam_121 + clinical covariates” and “Clinical covariates” models, and of **F)** “Cam_121” and “Clinical covariates” models. Out of a total of 194 patients, 23 were wrongly predicted by both the models in (E) and 22 were wrongly predicted by both the models in (F)

**Figure 1.**
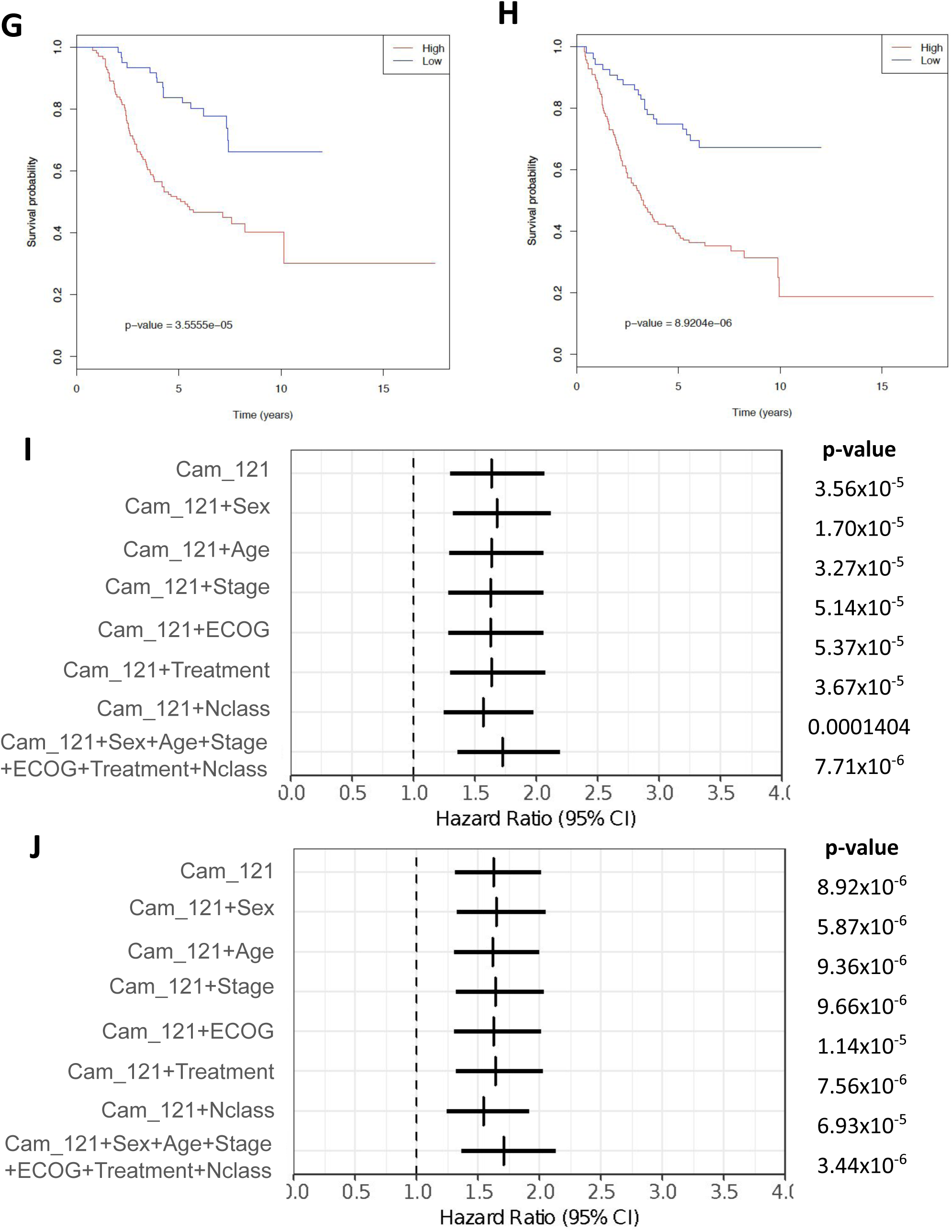
Cam_121 gene expression is strongly associated with both metastases and survival across multiple machine-learning and Cox regression analyses. Kaplan-Meier survival plots comparing the survival probabilities (y-axes) as a function of time (x-axes) of groups with high and low “Cam_121” (quantile 0.33 split) for **G)** overall survival (OS) and **H)** progression free survival (PFS). Forest plot indicating the hazard ratio estimates (vertical bars) and 95% confidence intervals (horizontal bars) related to the signature “Cam_121” when predicting **I)** OS and **J)** PFS by means of Cox proportional hazard models while controlling for different (sets of) clinical variables (y-axes). The Wald t-test p-value corresponding to the signature “Cam_121” parameter in each model is indicated on the right. ECOG; Eastern Cooperative Oncology Group Performance Status.

### 3. Signature added incremental prognostic value when combined with conventional clinical staging

In order to explore the relationship between Cam_121 gene expression and prognosis, we first performed univariate Cox regression using weighted Cam_121 expression score (see Methods) as predictor and found that it was significantly associated with both overall survival (OS; HR=1.64 (95% CI 1.30-2.07), p=3.56×10^−5^) and progression-free survival (PFS; HR=1.63 (95% CI 1.31-2.02), p=8.92×10^−6^) (Figure 1G-H). The Kaplan-Meier survival curves of melanoma patients with high and low weighted Cam_121 score (see Methods) showed a clear difference in survival between these two groups (Figure 1G-H). In order to evaluate whether the signature score contributed independent prognostic information while controlling for conventional clinical staging, multivariate Cox regression analyses were performed (Figure 1I-J). The signature score was significantly associated with both OS (HR=1.73 (95% CI 1.36-2.20), p=7.71×10^−6^) and PFS (HR=1.7 (95% CI 1.36-2.14), p=3.44×10^−6^) in multivariate Cox regression models.

We further tested the performance of Cam_121 in the (entirely separate) regional lymph-node metastasis samples within this dataset (n=177) and found that the weighted signature score was also significantly associated with both OS (HR=1.72 (95% CI 1.37-2.14), p=1.53×10^−6^) and PFS (HR=1.75 (95% CI 1.43-2.16), p=1.10×10^−7^) in multivariate Cox regression models (Supplementary figure S5), indicating that Cam_121 could also be relevant as a prognostic tool after the resection of stage III melanoma.

As Cam_121 was developed and tested on the same dataset, there are chances of overfitting. Therefore, we further tested the signatures’ prognostic power in an externally acquired independent validation dataset from the Leeds Melanoma Cohort (n=677) (15). This confirmed that Cam_121 was associated with melanoma-specific survival (MSS) in both univariate (HR=1.49 (95% CI 1.27-1.74), p=5×10^−7^) and multivariate Cox regression models (HR=1.7 (95% CI), p=0.001, Figure 2). A second external validation was attempted using the Lund primary melanoma cohort (20)(n=223), however only 24 of the Cam_121 genes were present in this dataset. This 24-gene signature was significantly correlated with PFS (HR=1.67 (95% CI 1.06-2.62), p=0.03 univariate Cox regression analyses), but not with OS (p=0.32), although there was paucity of signature genes within this dataset (Supplementary Figure S6).

**Figure 2.**
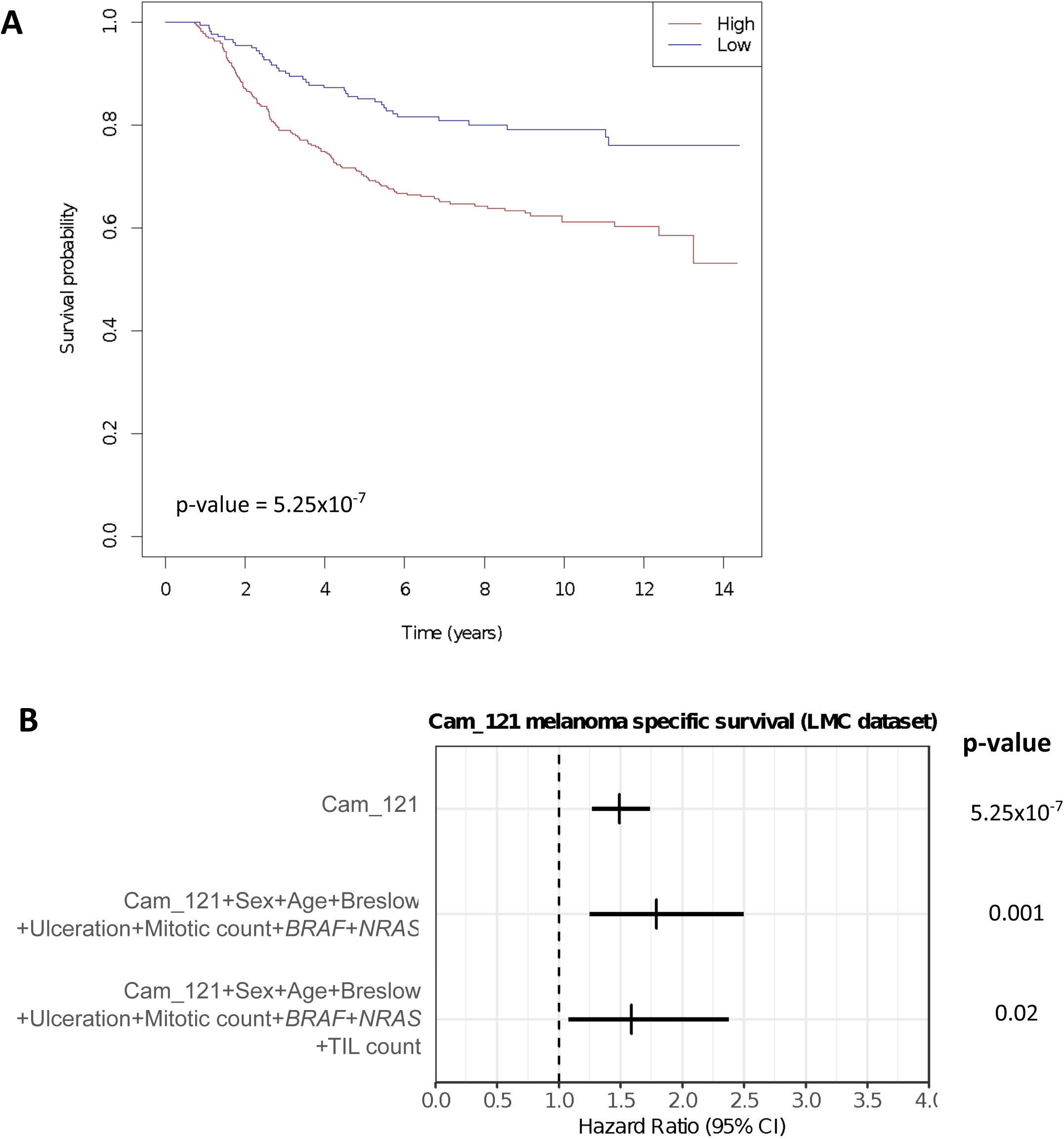
Validation of Cam_121 in an independently acquired external dataset (Leeds Melanoma Cohort, n=677). **A)** K-M curve comparing the Melanoma specific survival probabilities (y-axis) as a function of time (x-axis) of groups with high and low “Cam_121” (quantile 0.33 split) **B)** Forest plot indicating the hazard ratio estimates (vertical bars) and 95% confidence intervals (horizontal bars) related to the signature “Cam_121” when predicting Melanoma specific survival by means of Cox proportional hazard models while controlling for different (sets of) clinical variables (y-axes). Multivariate correction was undertaken for sex, age, breslow thickness, ulceration, mitotic count as well as *BRAF* and *NRAS* mutation status and (in the final row), correction was also undertaken for the tumour infiltrating lymphocyte (TIL) score. The Wald t-test p-values corresponding to the signature “Cam_121” parameter in each model are indicated on the right.

The published signatures from Thakur *et al* (16) (LMC_150; n=150 genes) as well as those from Gerami *et al* (12) (Decision-Dx Melanoma™; n=27 genes) were not found to be significantly associated with survival in multivariate models in our dataset, which may in part be reflective a higher proportion of stage III patients in these data than in these earlier studies (Supplementary figure S7).

### 4. Cam_121 gene signature score performed significantly better than genes selected at random in predicting overall and progression-free survival

In light of reports suggesting that randomly selected genes may perform equally well in predicting prognosis as published signatures (21), we decided to test the performance of Cam_121 against a set of 1000 randomly selected signatures of 121 genes each (from a pool of 19,434 protein-coding genes in our dataset; see Supplementary methods). We found that Cam_121 gene significantly outperformed random signatures across all measures of clinical efficacy including: OS (p=0.001) and PFS (p=0.001) in multivariate Cox regression models (Supplementary figure S8).

### 5. Stage II patients with a ‘high-risk’ signature demonstrated a 33% risk of death at 5-years, a threshold for which adjuvant therapy could be considered

We envisage that one of the central clinical applications of a prognostic GEP might be to identify those patients with stage II melanoma who may be at higher risk of relapse or death and for whom adjuvant systemic therapies may be considered. In order to compare our data with the registration adjuvant melanoma trials conducted in resected stage III melanoma survival (4-7), we measured the absolute risk of death at 5 years (calculated as the proportion of patients who died due to melanoma within 5-years from diagnosis).

Analyses in the Leeds Melanoma Cohort (where there was a higher preponderance of early-stage patients) revealed that stage II patients had a 27% (54/279) baseline absolute risk of death at 5-years. This risk rose to 33% (64/192) in those stage II patients with a high-risk weighted Cam_121 expression score profile and dropped to 14% (12/87) in those stage II patients with a low-risk profile (Table 1). The stratification of high/low risk cohorts in this context was based on a 0.33 quantile cut-off of the weighted Cam_121 expression score and subsequent references to high/low Cam_121 risk groups refer to these weighted expression groups (see methods).

**Table 1:**
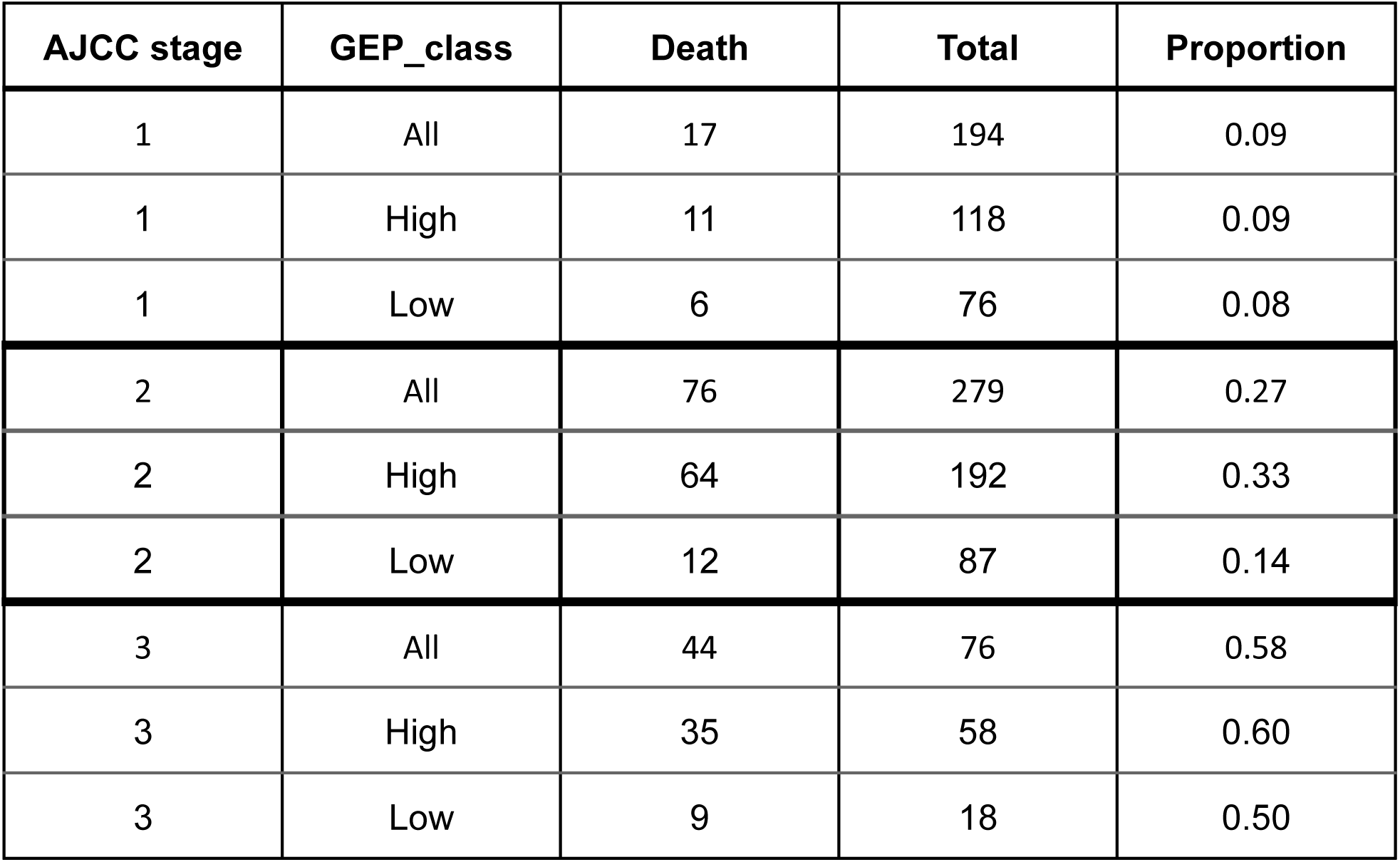
Clinical utility of GEP test 5-year melanoma specific survival (Leeds Melanoma Cohort Data). High and low risk expression cut-off are based on 0.33 quantile cut-off on weighted Cam_121 gene expression score expression of all 121 genes.

### 6. Per-gene analyses

In order to determine the relative influence of each gene and each baseline clinical covariate within the machine learning model, we analyzed their feature importance scores and found that no single feature dominated the performance of the model (Supplementary figure S9), suggesting that it is the combination of all the features that yielded improved performance over the baseline clinical covariates (Supplementary figure S4). In keeping with this, none of the Cam_121 genes proved significant in the per gene multivariate survival analyses after correcting for multiple testing (p-value<0.05), further confirming that it is the combination of all the genes that give improved performance in predicting OS and PFS (Supplementary figure S7).

We undertook further multivariate Cox regression analyses for all protein coding genes in this dataset (n=19,434) and found no single gene was significantly associated with either PFS or OS (Supplementary Tables S4 and S5).

### 7. A high weighted Cam_121 score reflected a lymphocytic depleted tumour with worse clinical outcomes

In order to identify the biological processes reflected by the signature, we inputted the genes into pre-ranked gene set enrichment analyses, ordered by their fold-change from the covariate-corrected differential expression analysis (see Methods). In doing this, we found that the top five significant (FDR corrected p-value<0.05) down-regulated hallmark gene-sets resulting from this analysis included interferon gamma (IFNγ) response, interferon alpha (IFN⍰) response, allograft rejection, inflammatory response and IL6-JAK-STAT3 signalling (Supplementary figure S11). Interestingly, when we ran gene-set enrichment on DEGs derived from the (entirely separate) lymph node samples (n=177; Supplementary figure S12), we observed significant (FDR corrected p-value<0.01) down-regulation of the exact same immune-related processes (Supplementary figure S12). Therefore indicating that the differential expression analyses (with a predominance of down-regulated genes in both the primary melanoma and regional lymph node datasets) also reflected a significant down-regulation of key immune mediated processes in the samples from patients that developed metastases (Supplementary figure S3 and S11).

We next used the Angelova dataset (22) to deconvolute the expression of immune cell subtypes within each sample (see Methods). We found a negative correlation between the weighted signature score and multiple immune cell types (with the highest correlation found with Activated B-cells (ρ=−0.75, p<2.2×10^−16^; Figure 3A), T-cells (ρ=−0.73, p<2.2×10^−16^; Figure 3B) as well the overall immune cell expression score (ρ=−0.73, p<2.2×10^−16^; Figure 3C). We also found that patients with samples showing a high weighted Cam_121 expression score were more likely to develop metastases than those with a low weighted Cam_121 expression score (Figure 3A-B & Supplementary table S6). Although 9 of the Cam_121 signature genes were common with the Angelova immune marker genes (*TUBB, AIM2, CASQ1, NTRK1, FASLG, CCR3, P2RY14, PRF1* and *CCR5*), we were able to demonstrate that samples segregated based on overall immune cell score, with low immune cell expression clustering with high weighted Cam_121 gene expression scores (Figure 3D), using principal component analysis. Therefore, we concluded that the weighted signature expression score negatively correlated with lymphocytic infiltration and that a high signature score equates to a relatively immune-deprived tumour with worse clinical outcomes.

**Figure 3.**
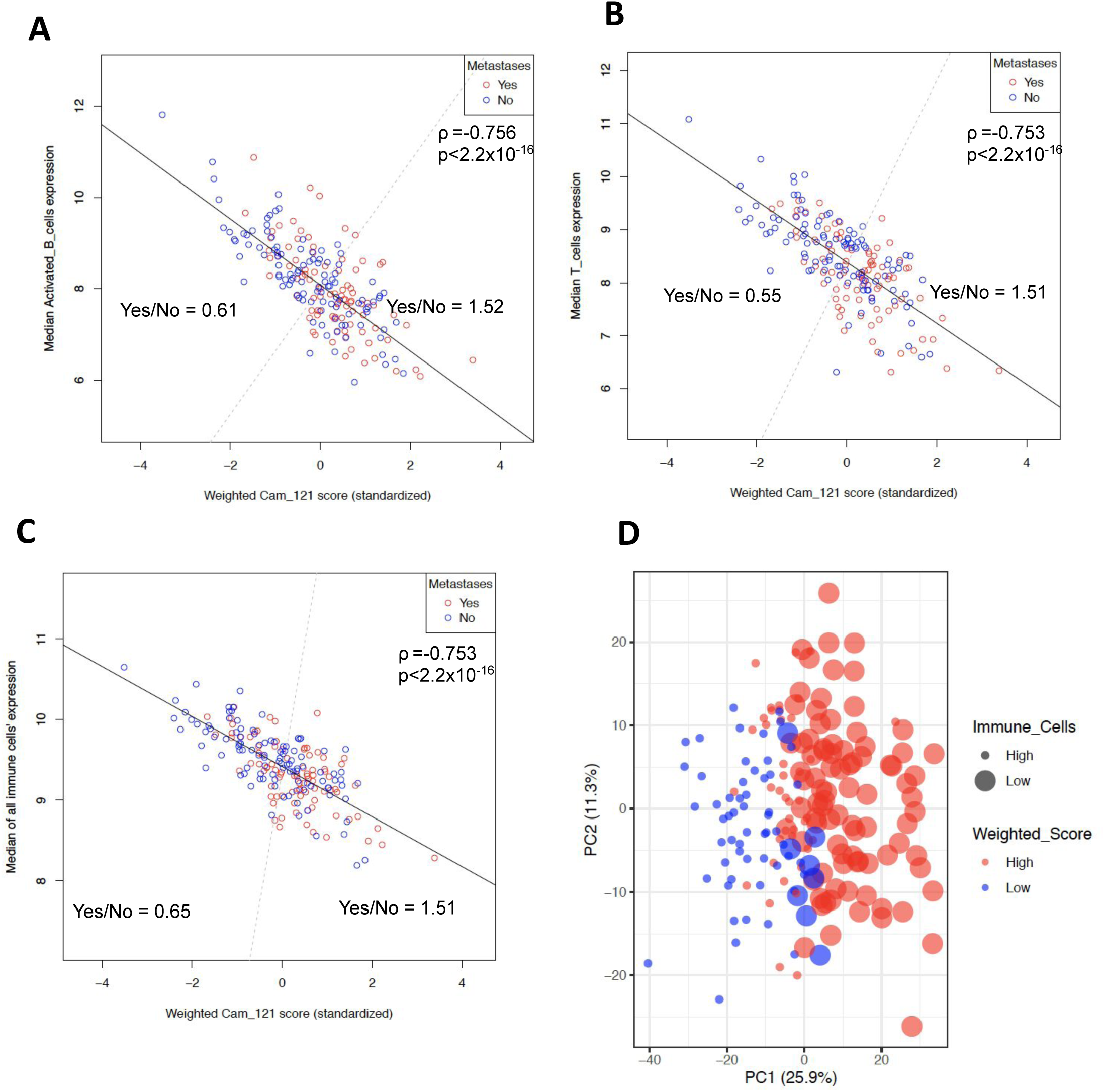
Weighted Cam_121 score negatively correlates with immune cell expression scores, indicating that a lower weighted signature expression score is associated with a richer immune microenvironment and better prognosis. Scatterplots showing the relationship between the standardised weighted Cam_121 score (x-axes) and **A)** the median Activated B-cell gene expression derived from Angelova et al, **B)** the median T-cell gene expression, and **C)** the total immune score (median gene expression of all cell-types in Angelova et al). Observations are colour-coded based on their metastatic status. A dashed grey separates all samples in two parts (passing through the median (x,y) coordinate). The Pearson’s correlation coefficient (ρ), its p-value and the ratio of metastases “Yes” to “No” samples on each side of the grey line is reported on the plot. **D)** Biplot of the scores of the observations on the two first dimensions of a PCA analysis of overall immune cell expression data. Observations are colour-coded according to their weighted Cam_121 expression score (“high”/”low” classification based on a quantile 0.33 split) and sized by overall median immune cell expression level (“high”/”low” classification based on a median split).

### 8. Gene reduction analysis revealed that Cam_121 had the potential to be reduced to upto 70 genes but was not prognostic in multivariate models

In order to assess whether the signature length could feasibly be reduced without comprising efficacy, we serially removed 10 genes at a time from Cam_121 ordered by decreasing fold-change (Supplementary Table 2; see Supplementary methods). In doing this we found that the top 70 genes with the highest fold-change (herein referred to as Cam_70) performed better in predicting metastases than signatures of all other lengths (Supplementary figure S13A-D). This reduced-length signature, Cam_70, further significantly correlated with both OS and PFS in uni- and multivariate survival models (p<0.05; Supplementary figure S13E-F) and was externally validated in the Leeds Melanoma Dataset (Supplementary figure S13G). However, the signature lost its significance after correcting for the tumour-infiltrating lymphocyte score in the multivariate model (p=0.14) and was therefore not considered for further analysis (Supplementary figure S13G).

## Discussion

The ability to identify primary melanoma patients at risk for disease recurrence is an important unmet need and effective prognostic biomarkers that could serve to guide adjuvant therapy are lacking. We sought to identify whether the expression of genes in a primary melanoma tumour can predict for distant metastasis and survival, analysing data acquired from one of the largest phase III prospective adjuvant melanoma clinical trials associated with long-term patient outcome data (18). We used covariate-corrected differential expression analyses to identify 121 genes significantly associated with distant metastases which made up our signature, and found that this added prognostic value in both the prediction of metastasis and survival. The prognostic relevance was further confirmed in an independent external validation cohort assessing melanoma-specific survival. Immune cell deconvolution analyses revealed that the weighted Cam_121 expression score negatively correlated with multiple measures of lymphocytic infiltration, with a high weighted signature score reflecting a relatively cold tumour-immune microenvironment with worse long-term prognosis. These findings were cross-validated using gene-set enrichment analyses, showing that differential expression analyses in both primary melanoma (n=194) and lymph node (n=177) datasets reflected down-regulation of the same key immune mediated processes. That this conclusion was reached using unbiased differential expression, re-affirms the central importance of the immune system in this setting.

The melanoma microenvironment consists of multiple immune and stromal cells, which play a critical role in regulating both the initiation and development of disease. Several studies have demonstrated the association of lymphocyte infiltration with longer survival (23-25), as well as an inverse relationship between tumour infiltrating lymphocyte (TIL) grade and the presence of lymph node metastases (26,27), implying that evaluating the tumour microenvironment landscape may hold promise for prognostic biomarkers. However only a limited number of studies have investigated the immune landscape in primary melanomas. A transcriptomic analysis of primary melanomas identified six distinct subgroups based on their expression of immune-related, keratin and beta-catenin pathway genes (28). In this study, patients with low immune but high beta-catenin score (CIC4) had the poorest overall survival (28). A recent study utilising high-throughput sequencing of T-cell receptor beta-chain in T2-T4 primary melanomas (n=199) indicated that the T-cell fraction accurately predicted progression-free survival and was independent of other key clinico-pathologic covariates (29). Although it was difficult to discern specific immune cell subtypes using bulk RNA sequencing, given that the weighted Cam_121 score was strongly negatively correlated with B cells, T cells and cytotoxic cells (ρ = −0.75, −0.73 and −0.74 respectively with p<2.2×10^−16^) and that interferon pathways dominated gene-set enrichment, we regard this as further evidence that a successful immune mediated cytotoxic anti-tumour response exists in primary melanoma. Critically, we found that the signature retained its prognostic power even after correcting for the TIL score, and it is our opinion that quantifying the expression of these key immune-mediated genes could potentially provide a more standardised and reproducible measure of immune activity. Furthermore, the prognostic relevance in both the primary melanoma and lymph node datasets attests to the signatures’ robustness. The challenge over the coming years will be to identify and validate a clinically-relevant measure of lymphocytic abundance of relevance to primary CM, that can be easily implemented in real-life clinical practice.

Interrogating the Leeds Melanoma Cohort, we found that the GEP-designated high/low risk could be used to separate patients with ≥ 33% risk of death at 5-years; a risk threshold for which we believe adjuvant systemic therapies could be considered. Conversely, it is envisaged that “low risk” GEP profiles could also be used to ‘downstage’ stage III patients for whom treatment might be unnecessary. There is substantial evidence supporting the importance of pre-treatment immune cell infiltration in eliciting anti-tumour responses with immunotherapy (30), however it remains to be seen whether Cam_121 expression can predict therapeutic responses in this setting. Future well-designed prospective clinical trials will ultimately be required to examine whether Cam_121 can be used to better tailor adjuvant therapy for early-stage melanoma patients.

The present study has a number of advantages over previous analyses. First, the large sample size linked to a well-conducted prospective clinical trial enabled an objective assessment of the risk of distant metastases, in addition to the key survival measures of interest. Furthermore, the long duration of follow-up (minimum of 6-years) in a cohort of patients predating modern approved adjuvant systemic therapies provided a unique insight into the ‘natural history’ of primary cutaneous melanoma. Finally, to our knowledge, this is the first large-scale biomarker analysis in primary melanoma to make use of data from comprehensive RNA sequencing. That such unbiased genome-wide assessment uncovered the dominance of immune-mediated genes reaffirms the central role of the host immune system’s ability to respond to the tumour resulting in immune editing or in some immune control.

It is important to point out that high quality evidence guiding the best practice use of gene expression predictors, particularly in the context of early stage CM are lacking. Future trials evaluating adjuvant therapies should examine both primary and locoregional melanoma samples using RNA sequencing technologies, to better characterise a molecular subtype/signature that could ultimately be used in conjunction with existing CM staging parameters and tailor future interventions more specifically to the individual. We believe that measures of lymphocytic infiltration should also be assessed in such studies.

Our results indicate that the Cam_121 signature score complements conventional melanoma staging by contributing prognostically relevant information and could potentially be used to select early-stage melanoma patients at higher risk of relapse or death. Further carefully designed prospective clinical trials will help guide how molecular features can be incorporated with traditional clinico-pathologic features to best estimate individual risk and guide the optimal clinical use of molecular biomarkers.

## Methods

### 1. AVAST-M melanoma cohort

This study made use of individual patient-level and transcriptomic data from the phase III adjuvant AVAST-M study, investigating the role of the angiogenesis inhibitor bevacizumab versus placebo in high-risk primary cutaneous melanoma (18). 1343 stage American Joint Committee on Cancer stage IIB (T3bN0M0 and T4aN0M0), IIC (T4bN0M0) and III (TxN1–3M0) cutaneous melanoma (7^th^ edition AJCC (19)) were recruited to the study over the period July 18, 2007-March 29, 2012, as previously described. The study (including the collection of DNA and RNA) was ethically approved in accordance with the Declaration of Helsinki (REC reference number 07/Q1606/15, 16^th^ March 2007). Participants provided written informed consented to sampling of their tumour blocks during study recruitment (and prior to the investigational systemic therapy).

All study participants underwent a sentinel lymph-node biopsy, and if positive proceeded to a completion lymph node clearance as per the study protocol. Demographic (including gender, age, centre, as well as pathologic data (site of primary, Breslow depth, ulceration, lymph node involvement and *BRAF*/*NRAS* mutation by pyrosequencing) was collected at the time of randomisation. Data was also collected on the timing, presence/absence and site/s of distant metastases (according to the findings from CT scanning). Data on overall and progression-free survival was collected with a minimum of 6 years follow-up.

RNA sequencing data was available on 204 primary melanoma samples and 177 regional lymph node (Ln) samples.

### 2. Leeds melanoma cohort

A primary melanoma transcriptomic dataset from the Leeds melanoma cohort study (LMC DASL array) was used as independent replication. This represents a population-controlled cohort study, as previously described (16). This study recorded data on melanoma-specific survival in 677 patients, calculated from the time of diagnosis to the time of last follow-up or time of death from melanoma, whichever occurred first. The regression coefficient (beta) for each gene (reflecting differential expression in AVAST-M dataset) was used to generate a weighted signature score in the new dataset. Hence for further analysis, a per-sample weighted gene expression score for our Cam_121 gene signature was calculated by multiplying the expression value of each gene by its corresponding beta coefficient (equation 1) followed by z-score normalization (zero mean-unit variance).

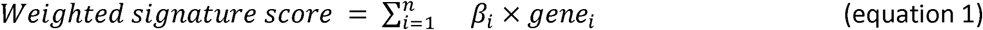

where i ranges from 1 to number of genes in the signature and β corresponds to the beta-coefficient of the respective gene obtained from DESeq2 analysis on AVAST-M melanoma cohort.

### 3. Lund melanoma cohort

Gene expression data on 223 primary tumors was generated using the Illumina DASL platform, as previously described (20). Data on relapse-free survival as well as overall survival were collected. The DASL platform analysed 7752 genes and only 24 of the Cam_121 genes were present. Validation was undertaken using weighted signature scores as outlined above.

### 4. mRNA extraction

Histopathological assessments of hematoxylin and eosin (H&E) stained slides were used to facilitate tumour sampling. Samples were consistently extracted from the least inflamed, least stromal regions of the invasive front of the tumour. RNA was extracted using the Roche High Pure FFPE RNA Micro Kit (cat# 04823125001) (Genentech Biosciences) according to the manufacturer’s recommendations. RNA quantity and quality were assessed using Agilent’s 2100 bioanalyzer.

### 5. Expression data generation

Extracted RNA was sequenced on the Illumina exome-capture sequencing platform, using 50 base-pair paired-end sequencing. Quality control (QC) was performed using fastq_utils (31) (version 0.14.7) and FastQC (32) (version 0.11.7). The reads that passed QC were aligned to the reference genome (GRCh38) using TopHat2 (33). Aligned reads were quantified using HTSeq (34). Only those genes with more than five reads, as reported by HTSeq, in at least one sample were selected for further analysis. The sequencing data was of good quality with a median of ∼50Million reads/sample. 446 tumor transcriptomes were profiled which included samples from: primary tumour (n=204); lymph node (n=177); local/distant relapse (n=58) and uncategorised samples (n=7). However, due to the clinical value of primary tumours in facilitating stratification at the earliest disease timepoint, we chose to focus our analyses on samples from cutaneous primaries (n=204) and used the lymph node samples as an internal validation.

### 6. Clinical covariate selection

Firstly, the association between distant relapse (yes/no) and clinical covariates was studied both with or without controlling for length of follow-up (defined as the time from diagnosis to last follow-up) and for treatment (yes/no). When ignoring length of follow-up and treatment, *generalised Cochran-Mantel-Haenszel tests* (R-package coin (35)) were used for nominal clinical predictors as they have the *Pearson’s Chi-square tests* and *Cochranl⍰-Armitage trend tests* as special cases when respectively considering the clinical covariate of interest as categorical or ordinal. For ordinal clinical covariates, we reported the ‘nominal/nominal’ association results when the ‘nominal/ordinal’ one was found less significant (as it is likely a sign that the assumption of linearity required by the ordinal test was not met. Mann–Whitney-Wilcoxon tests were used for continuous clinical covariates. When controlling for length of follow-up and treatment, likelihood ratio tests comparing the fits of logistic regression models with and without the clinical predictor of interest were used. The p-values were corrected for multiple testing using the Holm–Bonferroni method (Supplementary table S1A). Note that, as the 5-level Stage variable was highly related to N-class (Spearman correlation coefficient over 0.85), we picked the one with the lowest number of levels.

The variables 2-level Breslow staging and 2-level ECOG (Eastern Cooperative Oncology Group Performance Status) were significantly associated with relapse. The variable 2-level treatment was found to be related to relapse but is kept as control and the 2-level EventMet was the variable of interest indicating whether the patient relapsed or not. Therefore, the covariates Stage, Breslow thickness, ECOG and treatment were accounted for in the design formula of DESeq2 (36) without interactions and for further downstream analysis.

Secondly, the association between clinical covariates and overall survival (OS, calculated from the time of diagnosis to the time of last follow-up or death) and Progression free survival (PFS, calculated from the time of diagnosis to the time of last follow-up or death/progression to metastatic disease, whichever occurred first) was assessed by means of Cox proportional hazard models (R-package survival (37)). Both outcomes were considered as left-truncated due to delayed patient enrolment and right-censored due to loss of follow up or alive at the time of the end of the study. 6 years was chosen as the minimum cut-off for these analyses based on the original trial design.

Stage, Sex, Age and Nclass were significantly associated with both OS and PFS (p<0.05; Supplementary table S1(B-C)). The state of distant relapse (“EventMet”) was the most important variable but was not of our interest, hence dropped. ECOG is a good predictor. Treatment was not significant (p > 0.05) but was kept in the analysis. Also for PFS, ulceration (Ulc) was borderline at 5% level and was dropped from further analysis. Therefore, Stage, Sex, Age, Nclass, ECOG and Treatment were corrected for in subsequent gene-level survival analyses.

### 7. Differential expression analysis

Differential expression analysis between primary tumours that became metastastic vs those that remained non-metastatic over the six-year study period was performed using the package DESeq2 (36) (v1.18; R v3.6.1). The negative binomial models we considered controlled for the clinical covariates Stage, Breslow thickness, ECOG (Eastern Cooperative Oncology Group Performance Status) and treatment, as well as for the library size (offset). Raw read counts were provided as the input, with each column representing a sample and each row representing a gene, along with the categorical clinical information about each sample as colData. Samples with missing information for any of these four covariates were removed from the analysis, leaving 194 samples in total. The adjusted p-value cutoff (FDR) was set to 0.1 using the alpha parameter in DESeq2 *results* function and genes with FDR less than 0.1 were considered significantly differentially expressed.

Log fold change shrinkage was applied using the *lfcshrink* function with apeglm method from the apeglm package (38) (v1.6.0; R v3.6). For visualization and other downstream analysis, variance stabilizing transformation (vst) was used by means of the DESeq2’s *varianceStabilizingTransformation* function with option blind=FALSE.

### 8. Machine learning analysis

This section explains the steps followed to develop a machine learning classifier for each signature and to evaluate their performance in predicting relapse (yes/no). All the following steps were done using (R package caret (39) v6.0.84, DESeq2 v1.22.2; R v3.5.1)

#### 6.1 Dataset preparation and pre-processing

The dataset was prepared such that each column/feature contained information about all the patients/samples. These features can either be the expression values of the genes belonging to the signature of interest and/or the categorical clinical covariate information. In the latter case, the categorical clinical covariates were converted to numeric dummy variables. The clinical information about whether the patients metastasized or not were used as labels for the analysis.

This dataset was then divided into training and testing dataset for model selection and performance evaluation as explained below. In case of the gene expression data, vst transformation was applied to both training and testing dataset. This step was omitted for the clinical covariate dataset. To apply vst transformation on the testing dataset, mean-dispersion estimates learnt on the training dataset were used.

At each training step, the default parameters of the *trainControl* function (R-package caret) were used (i.e., freqCut=95/5, uniqueCut=10) with option search = “random”. The same feature(s) were removed from the testing dataset before evaluating the performance of the fully trained model.

#### 6.2 Model development and selection

For developing the machine learning classifier, nested leave-one-out cross-validation was performed for model development and evaluation:an inner LOOCV step first trained the model to prevent overfitting and the outer LOOCV loop then evaluated the performance of the model on the held-out sample (Supplementary figure S14). Therefore, at each step, 193 samples (i.e., n-1) were used for training the classifier using LOOCV whereas 1 sample was held-out for testing the performance of this final model. The process was repeated for all 194 samples, held-out one at a time and the final accuracy and kappa values were reported along with other performance metrics calculated using the *confusionMatrix* function (R-package caret).

Also, at each LOOCV training step, a random search was performed over 100 random combinations of hyperparameters and the set of hyperparameters giving maximum accuracy and kappa values on the training dataset was selected. This final model was then trained on the held-out sample. Note that nested-LOOCV was applied to compare the performance of different prognostic signatures on a per patient basis.

We selected all classifiers of the R-package caret that required less than 50,000 MB of memory, less than 5000 seconds of run time to execute this pipeline and did not produce “NA” as a possible metastases status. These eight selected classifiers are: Bayesian generalized linear model (40) (bayesglm), Lasso and elastic-net regularized generalized linear model (41) (glmnet), k-nearest neighbour (42) (knn), linear discriminant analysis (43) (lda), penalized logistic regression (44) (plr), random forest (45) (rf) and support vector machine (46) with linear (svmLinear) and radial kernel (svmRadial).

For each signature, each selected classifier was trained using the nested-LOOCV described above and the classifier giving the highest kappa value on the test dataset was selected as the best classifier for the signature of interest.

For the random forest classifier, the contribution of each feature (gene expression + clinical covariates) on predicting metastases, the feature importance score calculated for each LOOCV training step (1 step per held-out-sample; 194 in total) was saved and plotted as a boxplot using geom_boxplot function in R.

#### 6.3 Gene expression Vs covariate performance comparison per patient

In order to compare the performance of the signatures: Cam_121 + clinical covariates, Cam_121 only and clinical covariates only on a per patient basis, the test prediction of the best performing classifier (random forest (rf), svmLinear and glmnet respectively) for each signature was extracted. To visualize the overlap, venn diagram was generated using the *venn.diagram* function from the VennDiagram package (47) (1.6.20) in R.

#### 6.4 Statistical analyses

To check if the performance of various gene expression signatures in predicting relapse (yes/no) across multiple classifiers (n=8) is significantly better than those built on clinical covariates alone, Welch Two Sample t-tests were used (R function *t.test* function with option var.equal=FALSE, paired=FALSE, alternative=“greater”) for each performance metric and each combination of signature and clinical covariate at the 5% level. The null hypothesis was that the true difference in mean performance across 8 classifiers between both conditions (‘clinical covariates alone’ versus ‘clinical covariates and signature’) equals 0 while the alternative hypothesis was that the true difference in means is greater than 0.

### 9. Determination of the weighted expression score cut-off to define ‘high’ and ‘low’ absolute risk of death at 5-years

Data from the Leeds Melanoma Cohort was used to calculate the absolute risk of death at 5-years (this dataset was chosen for this analysis due to the preponderance of early stage patients; Stage I = 194 samples; Stage II = 279 samples; Stage III = 76 samples). Five-year melanoma specific survival was calculated such that those patients who died due to melanoma within 5 years of follow-up were assigned event = “Yes” and those that did-not were assigned event = “No”. Those patients who did-not yet die and were followed up for less than 5 years were removed from the analysis due to inadequate follow-up.

The quantile cut-offs 0.25, 0.33 and 0.5 were used to divide patients into high/low groups based on their corresponding weighted Cam_121 expression score. Absolute risk of death at 5-years was calculated as the ratio of patients where event = “Yes” to the total number of patients within each stage (I-III). The cut-off giving the maximal separation (of absolute risk of death) between high/low groups was selected. This was achieved using a 0.33 quantile cut-off of the weighed Cam_121 expression score and subsequent references to high/low Cam_121 risk groups refer to these high/low weighted stratification cohorts.

### 10. Survival analyses

For each sample, a vector of weighted signature expression scores was calculated by using equation 1 on the vst normalized gene expression data. The standardised scores were then used as a continuous predictor in Cox regression models fitted by means of the coxph function of the survival package (37) (c2.42.3) in R (v3.5.1). The hazard ratio (HR) (95% CI) and p-values corresponding to the signature were reported in both univariate and multivariate analyses.

In order to display Kaplan-Meier survival curves, samples were divided into “high”/”low” signature expression groups based on the 0.33 quantile cutoff which we obtained from the absolute 5y risk assessment in Methods section 8. Samples with weighted signature expression score greater than this cutoff were assigned to the “high” group and those with weighted signature expression score lower than this cut-off were assigned to the “low” group. The survival distribution of both groups were finally compared by means of Log-rank tests using *survfit* function from the survival package (v2.42.3) in R (v3.5.1)).

### 11. Tumour immune microenvironment analysis

Sample-level gene expression data from the AVST-M cohort was deconvoluted into infiltrating immune cell scores using the Angelova dataset (22). This dataset reports 812 marker genes corresponding to 31 immune cell-subtypes. Out of these 812 genes, 53 genes were missing from our 38,690 gene list. Therefore, two immune cell subtypes MDSC (myeloid-derived suppressor cells) and NK56_bright (natural-killer CD56^bright^ cells), with more than 1% of missing marker genes were removed from further analysis, leaving 719 marker genes corresponding to 29 cell-types.

#### 11.1 Immune cell correlation analysis

To perform correlation analysis for each cell type, the median of the corresponding marker genes’ expression for each sample (y-axis) was plotted against the weighted Cam_121 expression score for that sample (x-axis). To calculate the overall immune score, the median of all 719 marker genes was used for the analysis. A regression line was fitted through these points using the *lm* function of the stats package (48) (v3.6.2) in R (v3.6.2). The Pearson’s correlation coefficient (ρ) between the immune-cell score and the signature, as well as the p-value of the corresponding test of association were estimated by means of the function cor.test of the stats package (v3.6.2) in R (v3.6.2). Sensitivity analyses considering robust regression (*lmrob* function of the robustbase package (49) (v0.93.5)) led to similar conclusions.

Samples were coloured according to their metastatic status. With red points indicating the samples obtained from patients who later metastasized (*Yes*) and blue points indicating the samples obtained from patients who didn’t (*No*). To calculate the distribution of these 194 samples along the axes, they were equally divided into two parts of 97 samples each and the ratio of metastases vs non-metastases samples samples was calculated for each part using equation 2. To equally divide these samples, a perpendicular line was drawn to the fitted line obtained above which passed through the median (x,y) point.

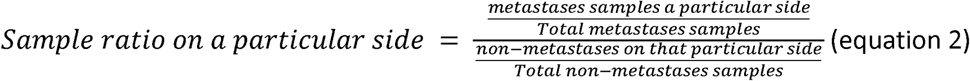

To confirm the relationship between the signature and immune cells, a PCA analysis was performed. For each cell-type, the corresponding marker genes’ expression was projected into the principal component space and the first two principal components explaining the maximum variance were plotted against each other. These samples were then colored by the “high”/”low” weighted Cam_121 expression score groups obtained during the survival analysis and sized by the “high”/”low” groups based on marker genes’ median expression corresponding to that particular cell-type. Here, to divide the samples into two independent categories based on their marker genes’ expression, the median cut-off was used, where the sample with marker genes’ median expression value above its overall median value was assigned to the “high” group and that with marker genes’ median expression value below its overall median value was assigned to the “low” group.

#### 11.2 Gene set enrichment analyses

Preranked GSEA (GSEA-P) was implemented using the *GSEAPreranked* tool of the GSEA software from Broad Institute (50,51) (v4.0.2). Hallmark gene sets were downloaded from the MSigDB database (52) (6.2.0). The genes were pre-ranked according to their shrinked log fold change values obtained in the differential expression analysis and the GSEAPreranked tool was run with default parameters with the enrichment statistic set to “classic”.

## Data availability

All the raw RNA sequencing data (forward and reverse fastq files) will be made available at the European Genome-Phenome Archive (https://www.ebi.ac.uk/ega/ at the EBI).

## Funding

This work was supported by Cancer Research UK and the Wellcome Trust. Bevacizumab was supplied by Genentech pharmaceuticals. NAF was partially supported via the European Union’s Horizon 2020 research and innovation programme under grant agreement No 668981. The Leeds Melanoma Cohort research was carried out with research funding CR-UK; C588/A19167, C8216/A6129 and C588/A10721 and NIH; CA83115. MRM is supported by the National Institute for Health Research (NIHR) Oxford Biomedical Research Centre. The views expressed are those of the authors and not necessarily those of the NHS, the NIHR or the Department of Health.

## Acknowledgements

We would like to thank Neera Maroo, Leticia Campo and the Translational Histopathology Laboratory at the Oxford Department of Oncology for sample biobanking and processing. Vivek Iyer for cross-checking the differential expression analyses. Wolfgang Huber and Janet M. Thornton for advice on machine learning analyses. Michael I. Love for advice on data normalization for machine learning. Danish Memon for expert input on immune cell deconvolution and Andrea Manrique-Rincon for her help on the critical interpretation of these analyses. Alastair Droop for critical review of the manuscript.

## Author contributions

RR, AB, and DJA conceived the project. MRM co-ordinated the sample biobanking. MW and YY co-ordinated the mRNA extraction and RNA sequencing. RR co-ordinated the sequencing and clinical data extraction. NAF derived the RNA-seq counts from raw sequencing reads. RR, DJA, AB and MG developed the clinical questions and experiments and MG carried out the analysis. DLC ran the statistical analyses. JN, TB and JNB validated the findings in the Leeds Melanoma Cohort. ML and GBJ validated the findings in the Lund Melanoma Cohort. PC, MRM and CP provided senior input on the translational scope of the project. MG and RR wrote the manuscript. AB and DJA provided overall study supervision. All authors approved the final manuscript.

---

**Supplementary Figure S1.**
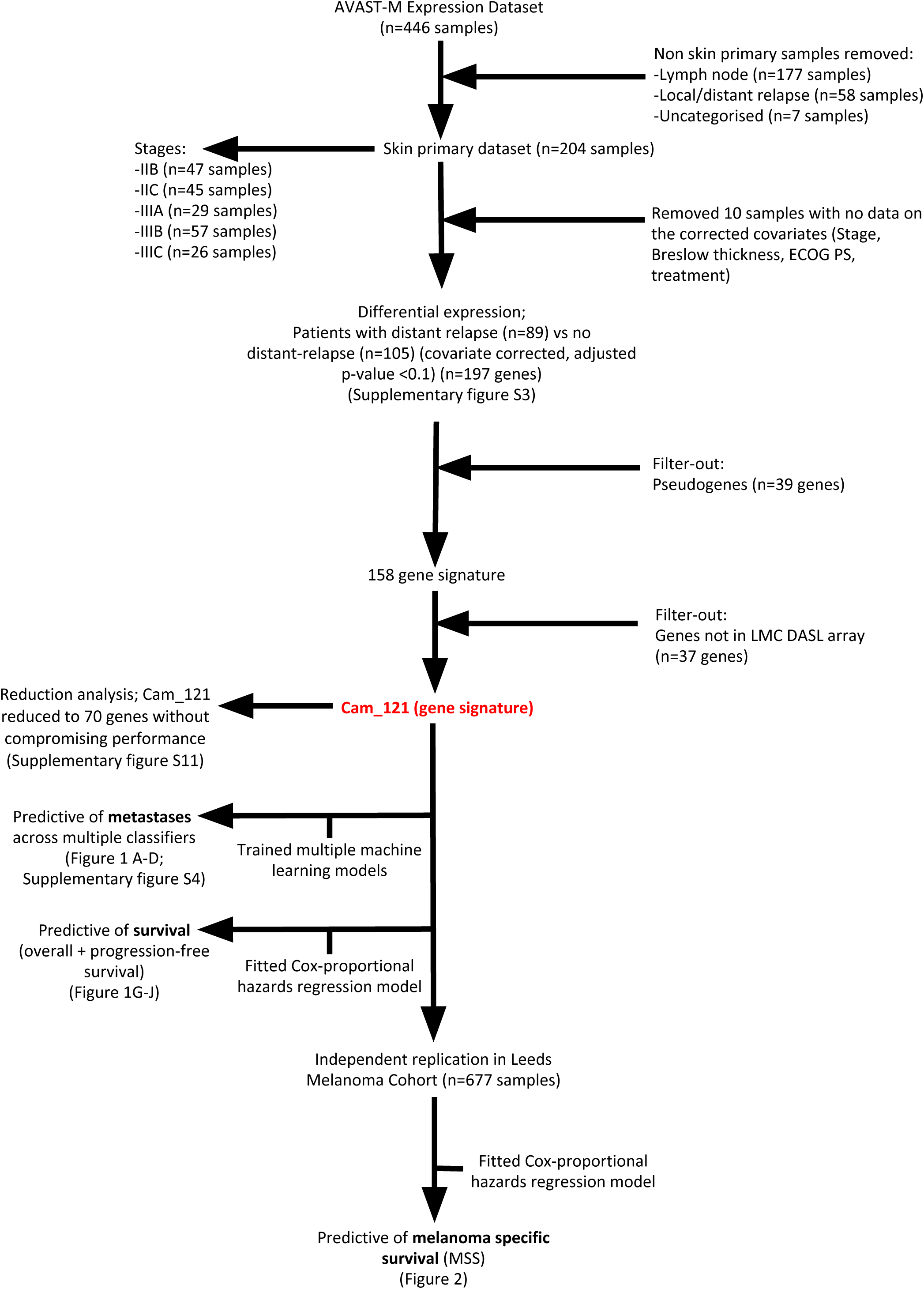
Flow chart of the analysis. The figures corresponding to each analyses are indicated in bold.

**Supplementary Figure S2.**
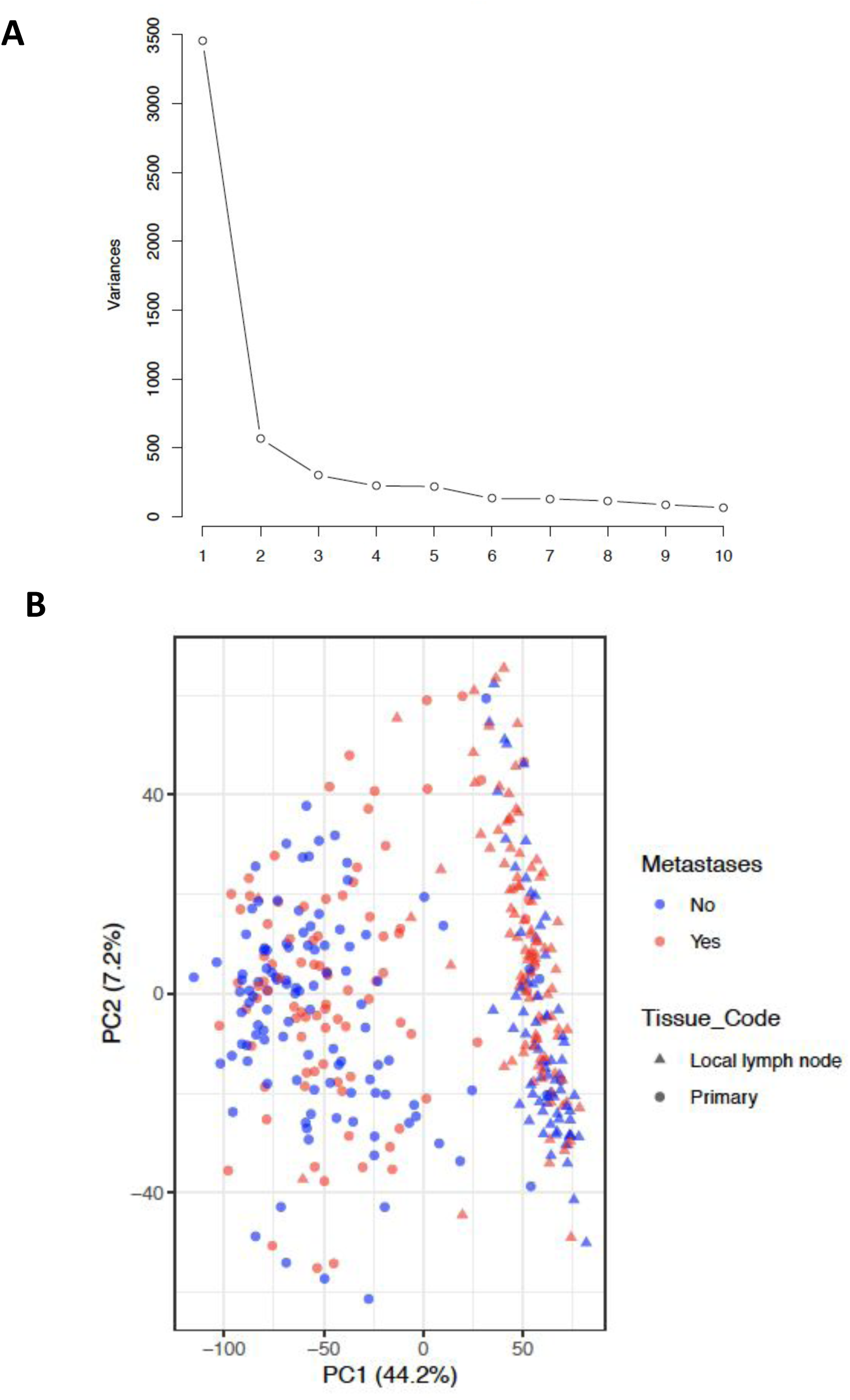
PCA results of the 1000 most variable genes, showing that samples separated by their site of origin. **A)** Scree-plot showing the variance of the data (y-axis) explained by each of the first 10 principal components (x-axis). **B)** Biplot of the scores of the observations on the two first dimensions of a PCA analysis of expression values of 1000 most variable genes. The samples are colour-coded according to their relapse status (yes/no) and shaped according to their tissue of origin (skin samples in circles, lymph nodes in triangles). The plot shows that samples cluster based primarily on the tissue of origin, separated primarily on the x-axis accounting 44% of the total variance.

**Supplementary Figure S3.**
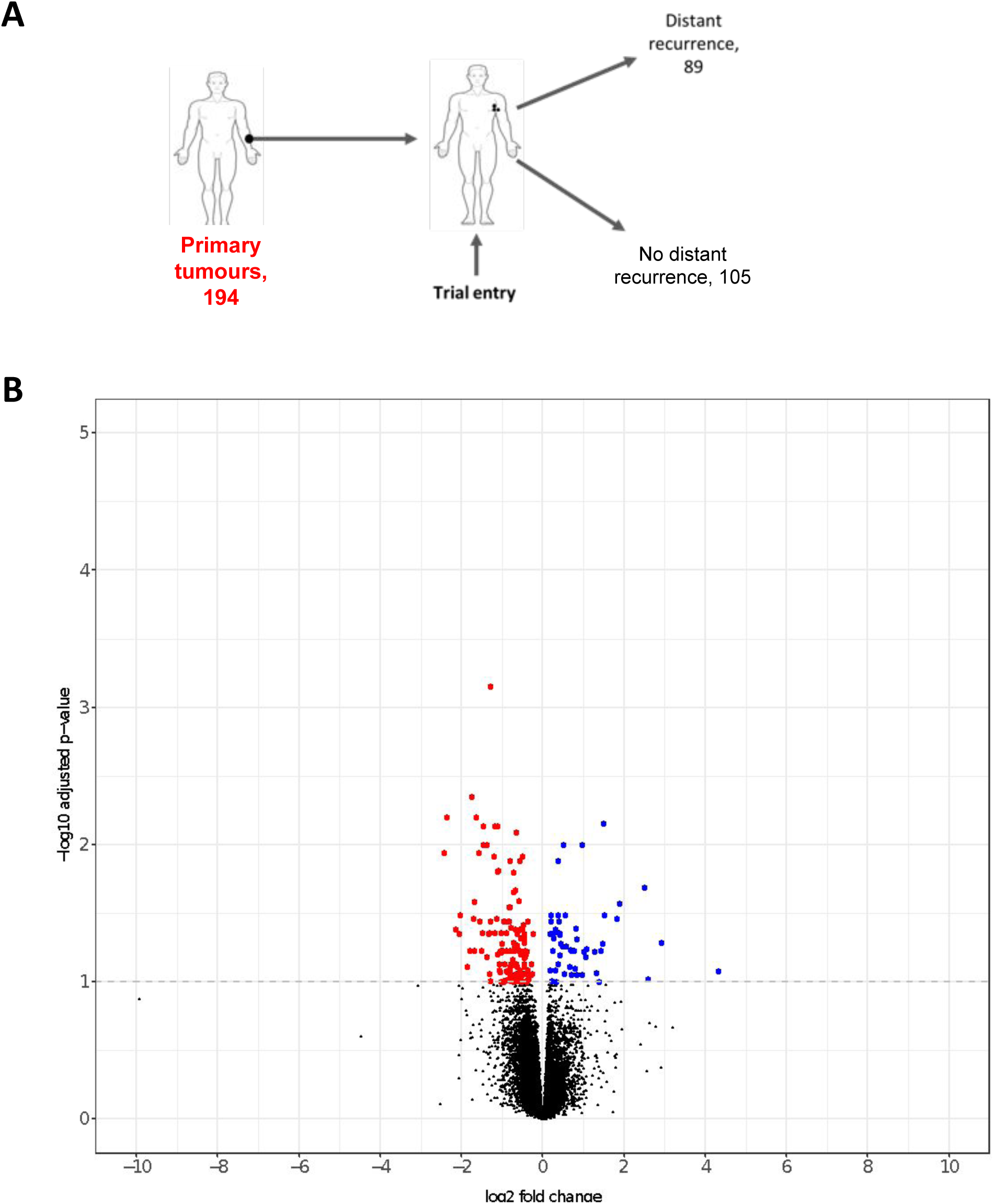
Results of metastases vs non-metastases covariate-corrected differential expression analysis for primary melanoma samples, indicating predominantly down-regulated DEGs. **A)** Schematic of the number of samples used in the differential expression analysis. **B)** Volcano plot showing, for each gene, the negative of the log 10 of its FDR corrected p-value (y-axis, FDR<0.1) and its log 2 fold change estimate according to the differential expression analysis results. The downregulated DEGs are coloured red (144/197, 73.1%) and the upregulated DEGs (53/197, 26.9%) are coloured blue.

**Supplementary Figure S4.**
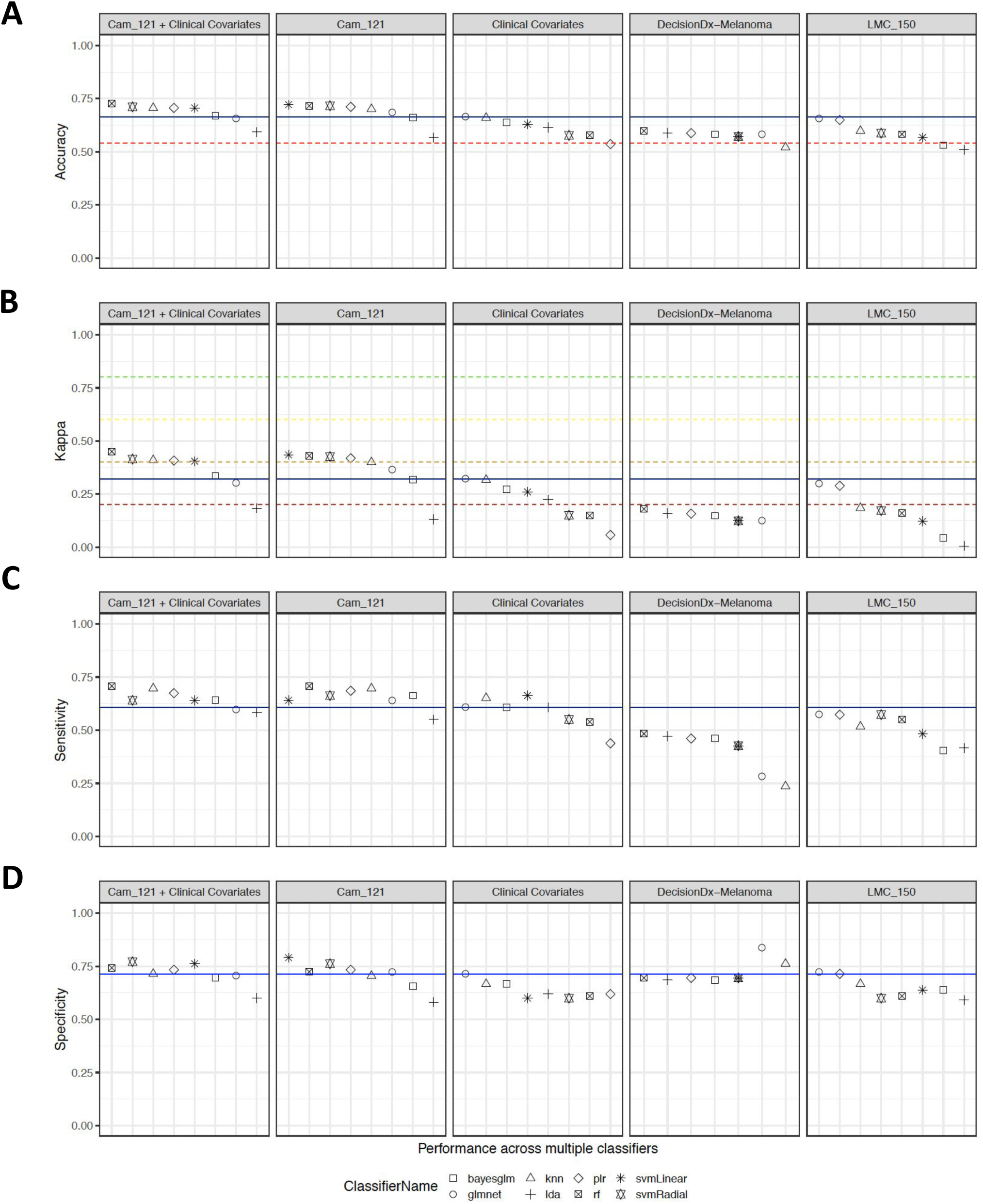
Machine learning results for multiple classifiers showing that Cam_121 outperforms clinical covariates as well as published prognostic signatures. Scatter plots showing, for each set of predictors (columns), the scores of **A)** accuracy **B)** kappa **C)** sensitivity **D)** and specificity (y-axes) of the 8 selected classifiers, ordered by performance on predicting accuracy from top to low (x-axes). To facilitate comparisons, a blue line representing the performance of clinical covariates based on the glmnet classifier (which was found to be the best performing classifier for this set of predictors for all criteria except sensitivity) was added to each plot The red line in (A) represents the accuracy achieved when the classifier only predicts the majority class (metastases=“No” in this case 105/194 (54.12%)). Coloured lines in (B) represent the range of agreement bands of [CITE WHOEVER] defined for Cohen’s Kappa values. Refer to supplementary table 3 for a statistical comparison between these signatures. Decision-Dx Melanoma: Decision-Dx Melanoma™, LMC_150: Leeds Melanoma Cohort 150 gene signature.

**Supplementary Figure S5.**
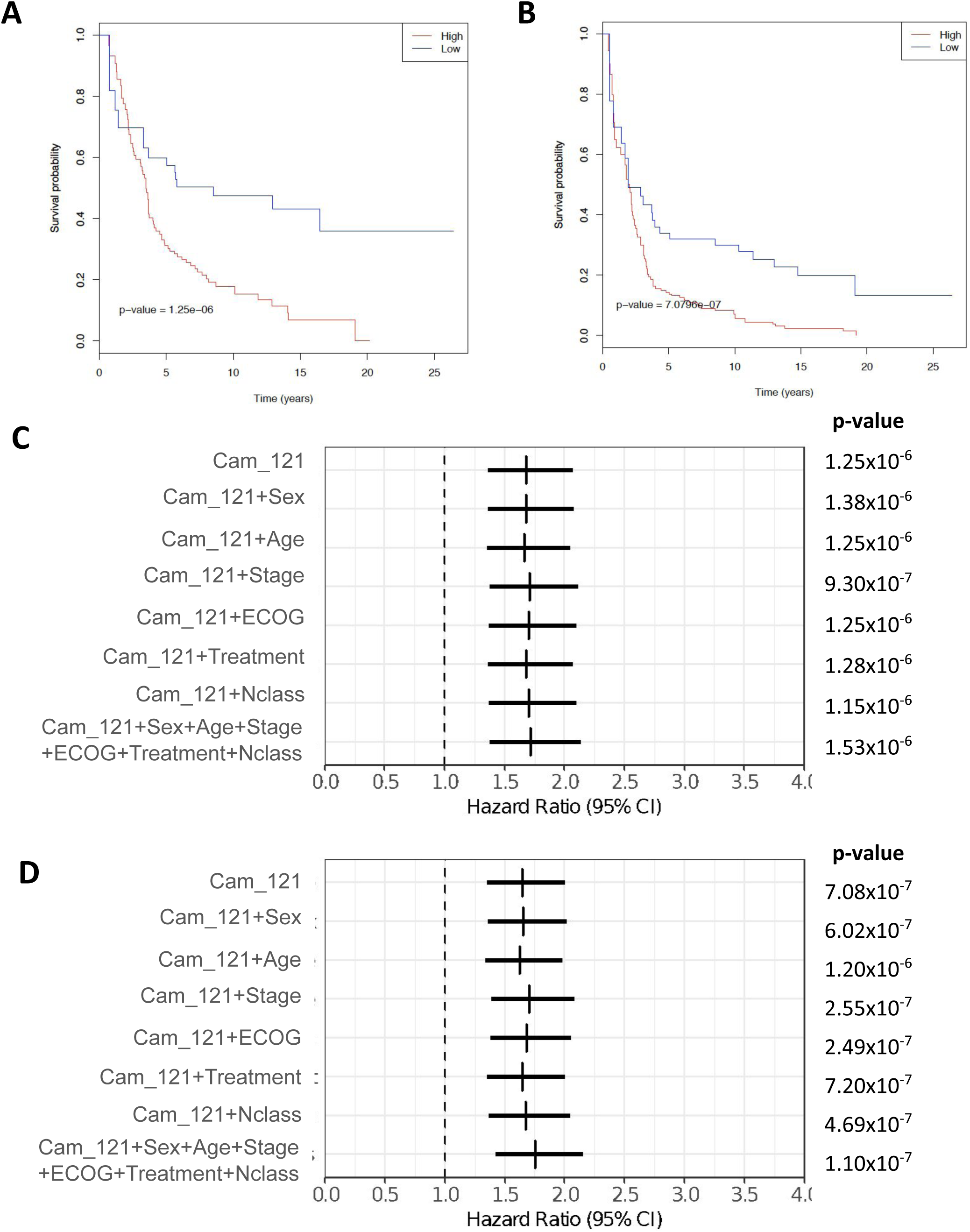
Validation of Cam_121 in the entirely separate lymph node samples from the AVAST-M cohort (n=177). K-M curve comparing the **A)** OS probabilities (y-axis) and **B)** PFS probabilities (y-axis) as a function of time (x-axis) of groups with high and low absolute 5y risk of death (0.33 quantile split). **C)** Forest plot indicating the hazard ratio estimates (vertical bars) and 95% confidence intervals (horizontal bars) related to the signature “Cam_121” when predicting OS and **D)** PFS by means of Cox proportional hazard models while controlling for different (sets of) clinical variables (y-axes). The Wald t-test p-values corresponding to the signature “Cam_121” parameter in each model are indicated on the right.

**Supplementary Figure S6.**
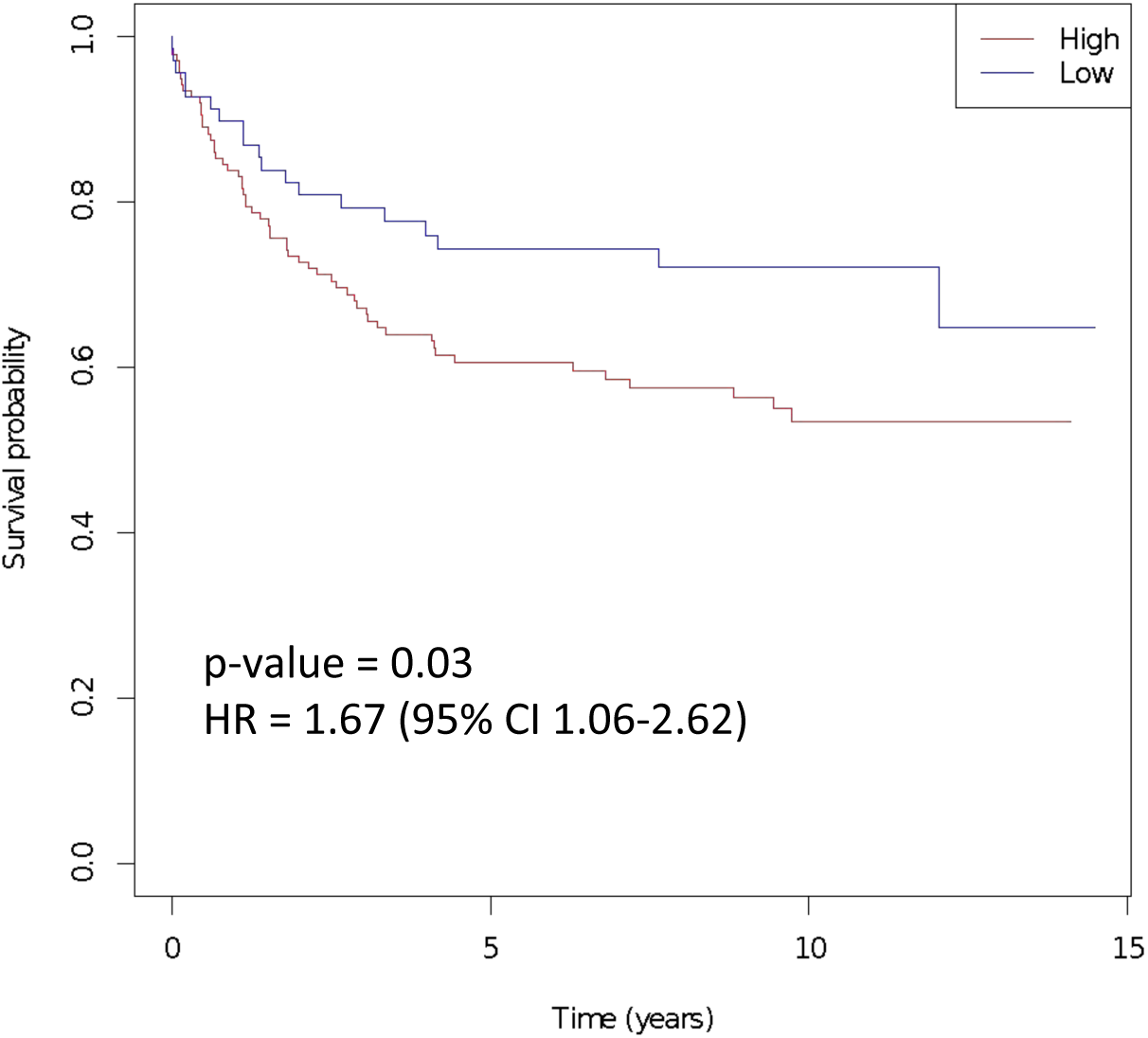
Validation of Cam_121 within a second independently acquired external dataset (Lund Melanoma Cohort, n=223). **A)** K-M curve comparing the progression free survival probabilities (y-axis) as a function of time (x-axis) of groups with high and low “Cam_121” (quantile 0.33 split).

**Supplementary Figure S7.**
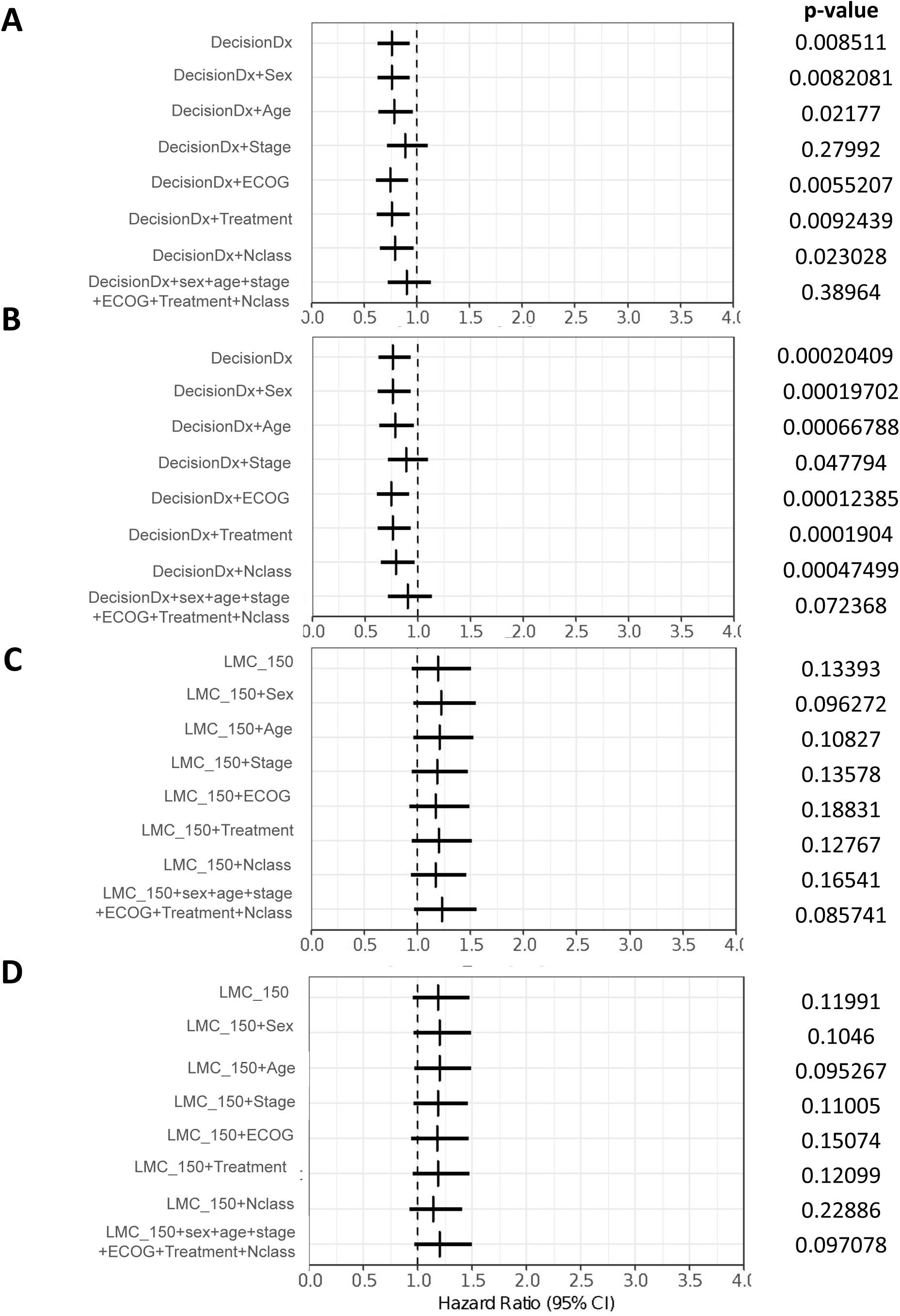
Forest plot of univariate and multivariate survival analysis for previously published signatures (Gerami and LMC). Forest plot indicating the hazard ratio estimates (vertical bars) and 95% confidence intervals (horizontal bars) related to the signatures of Gerami (**A** and **B**) and LMC (**C** and **D**) when predicting overall survival (**A** and **C**) and progression free survival (**B** and **D**) by means of Cox proportional hazard models while controlling for different (sets of) clinical variables (y-axes). The Wald t-test p-values corresponding to the parameter of the signature of interest in each model are indicated on the right. ECOG; Eastern Cooperative Oncology Group Performance Status.

**Supplementary Figure S8.**
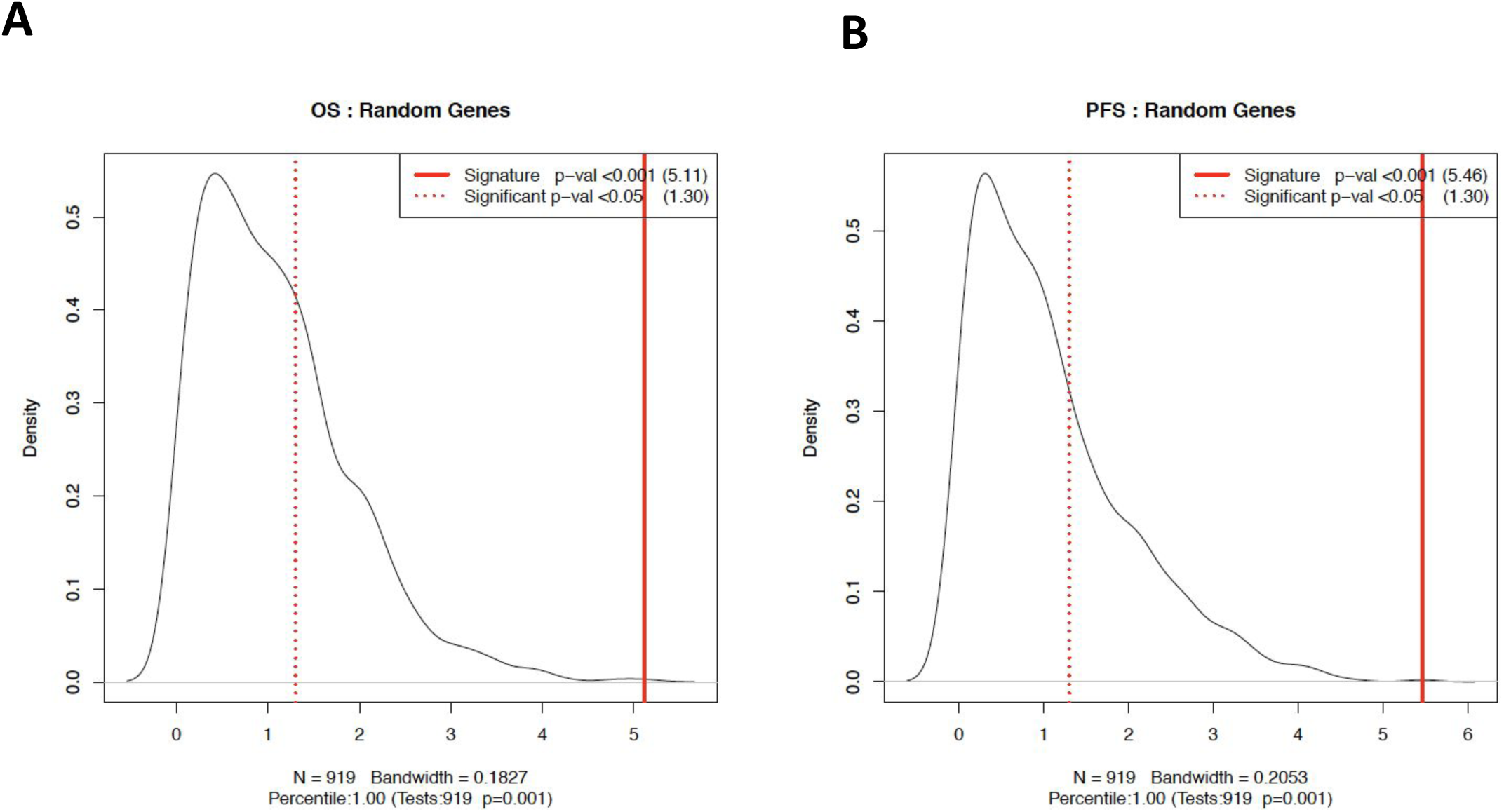
Analysis of Cam_121 showing that the signature significantly outperforms 1000 sets of 121 randomly selected protein-coding genes. Comparison of the performance of the Cam_121 gene signature (red vertical lines) and random signatures (smoothed densities) in terms of **A)** multivariate OS and **B)** multivariate PFS. The p-values are defined as the fraction of scores of random signatures which are greater that the one observed when using the (real) Cam_121 gene signature. The red vertical lines correspond to the cut-off of 5% one-sided tests.

**Supplementary Figure S9.**
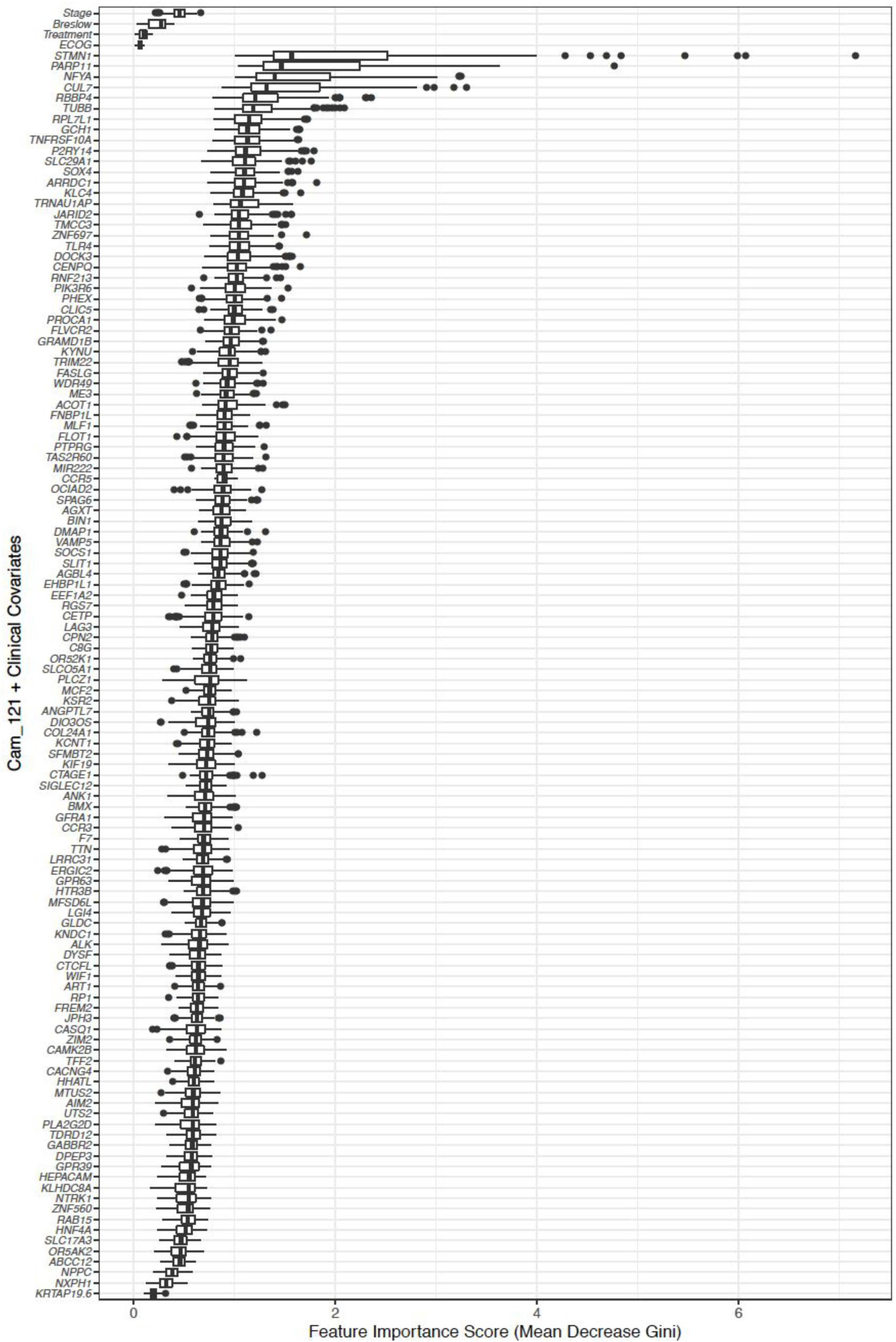
Feature importance score (FIC) for each gene in Cam_121 showing similar scores across all genes, thereby suggesting that it is a composite analysis of all genes that contribute to the signatures’ prognostic power rather than the dominance of any single gene. Genes are ranked in decreasing FIC order. The clinical covariates are indicated on the top for comparison. Each boxplot highlights five summary statistics including median, first and third quartiles, two whiskers and all the outlying points. More information can be found on the official documentation of the *geom_boxplot* function.

**Supplementary Figure S10.**
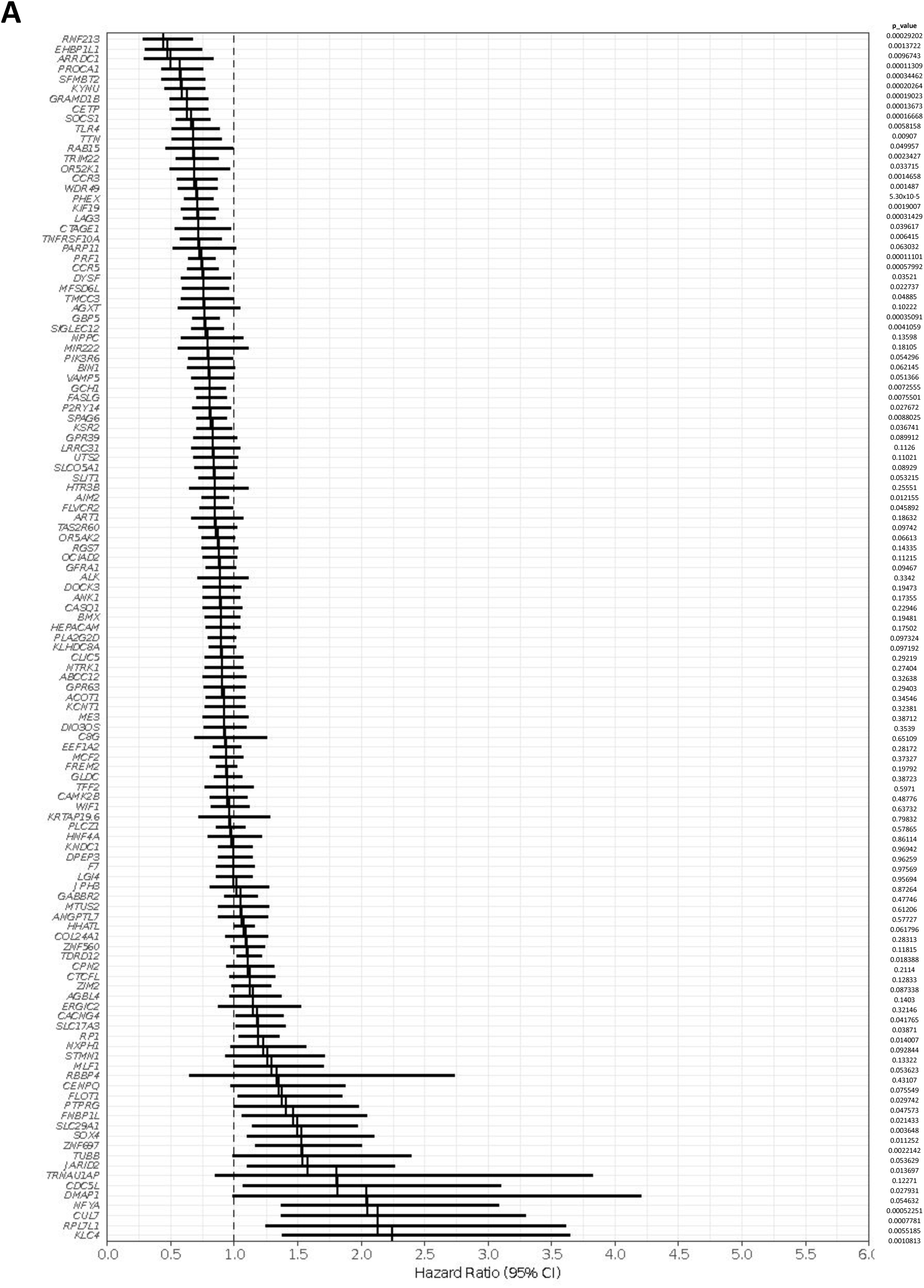
Multivariate survival analysis for each gene in Cam_121 showing some genes can significantly predict survival, however none proved significant after correction for multiple testing (FDR corrected p-value<0.05). This therefore suggests that it is a composite analysis of all genes that contribute to the signatures’ prognostic power rather than the dominance of any single gene. **A)** OS. Genes are ranked in increasing order of HR. The Wald t-test p-values (before FDR correction) corresponding to the parameter of the signature of interest in each model are indicated on the right.

**Supplementary Figure S10.**
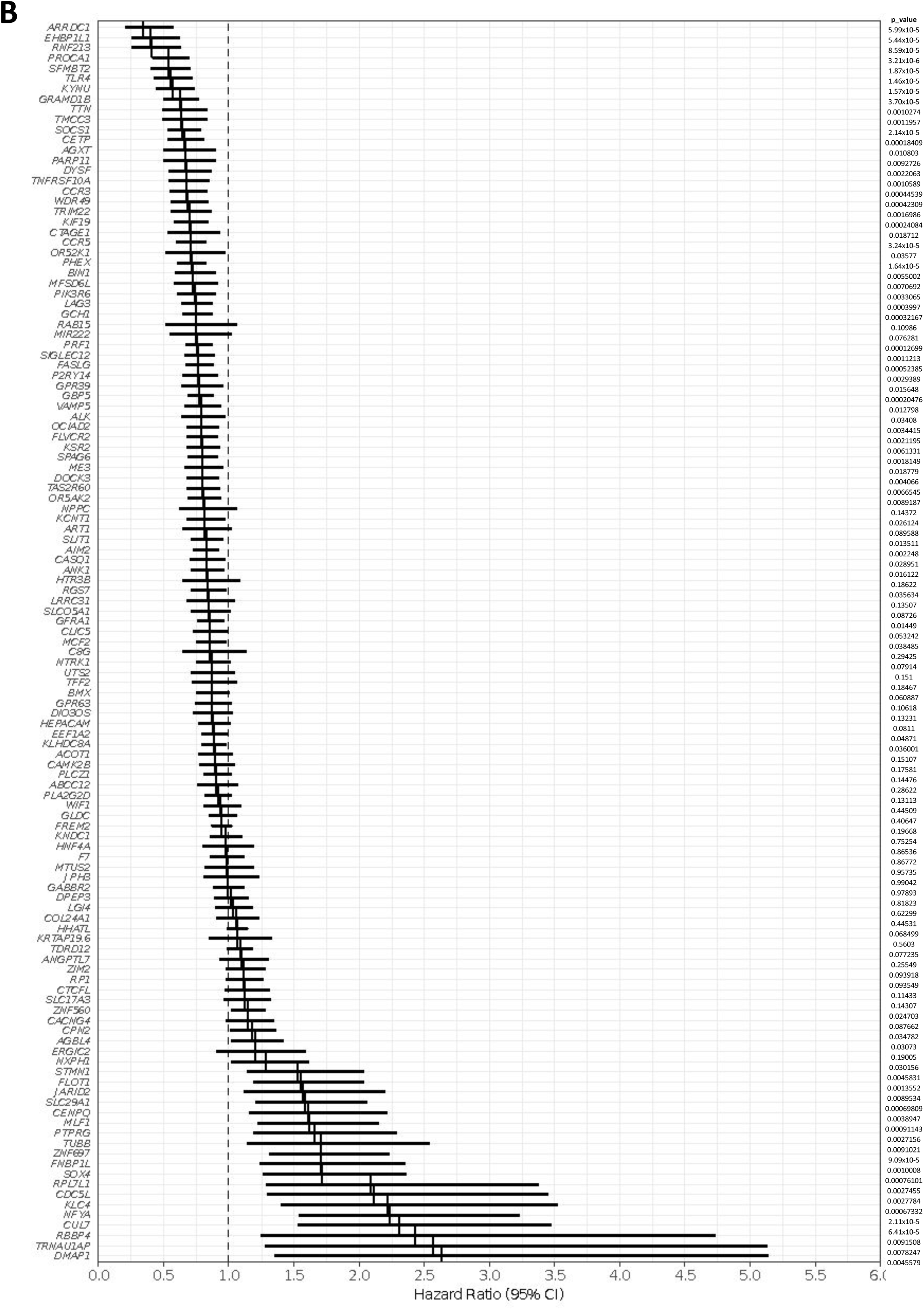
Multivariate survival analysis for each gene in Cam_121 showing some genes can significantly predict survival, however none proved significant after correction for multiple testing (FDR corrected p-value<0.05). This therefore suggests that it is a composite analysis of all genes that contribute to the signatures’ prognostic power rather than the dominance of any single gene. **B)** PFS. Genes are ranked in increasing order of HR. The Wald t-test p-values (before FDR correction) corresponding to the parameter of the signature of interest in each model are indicated on the right.

**Supplementary Figure S11.**
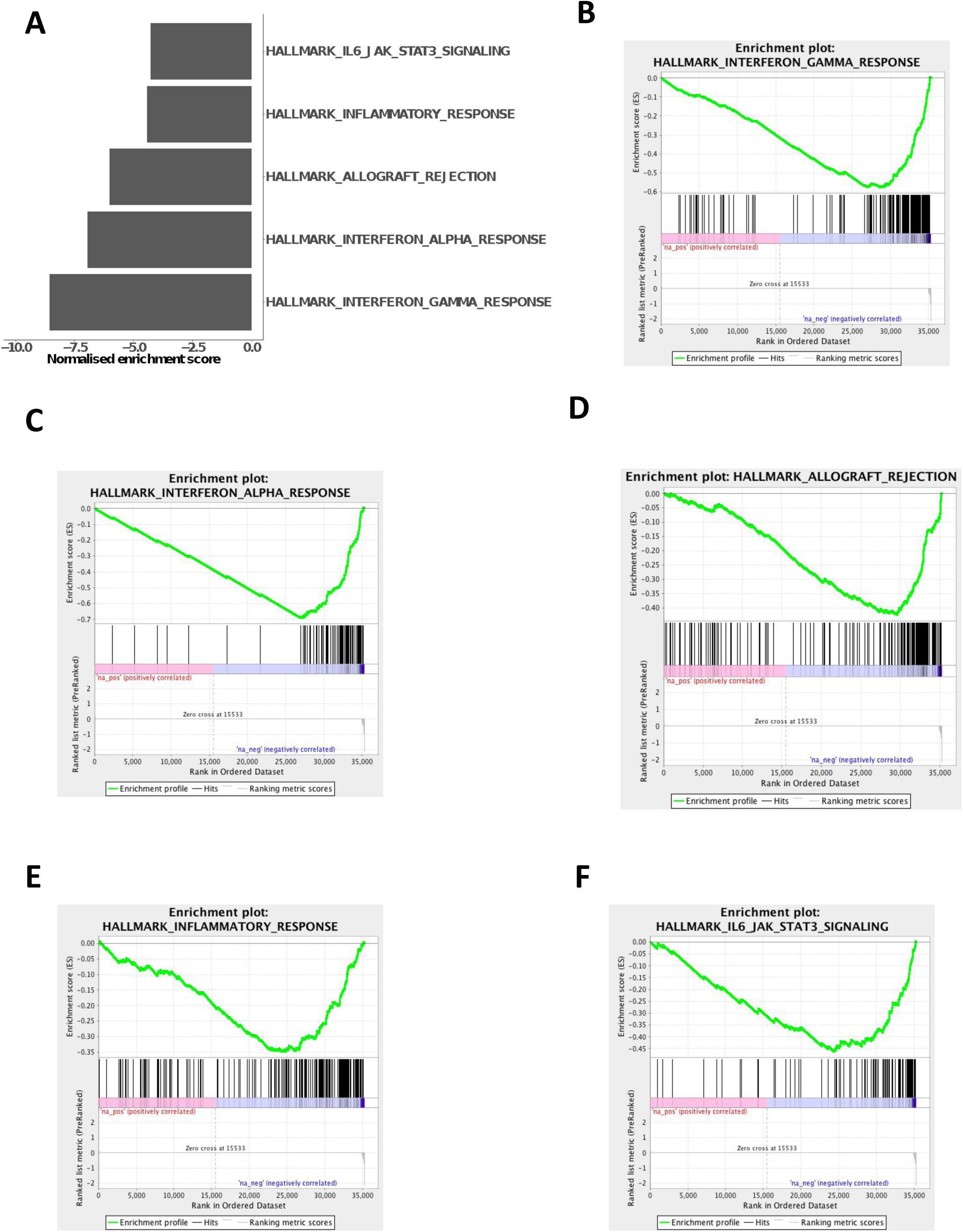
Results of gene set enrichment analysis showing downregulation of key immune-related pathways in skin samples. A) Barchart of the top five most significant down-regulated Hallmark gene sets (p<0.01 for all gene sets) B-F) The corresponding enrichment plots from (A). The top portion of the plot shows the running enrichment score for the gene set as the analysis walks down the ranked list. The middle portion of the plot shows where the members of the gene set appear in the ranked list of genes and the bottom portion of the plot shows the value of the ranking metric as one moves down the list of ranked genes.

**Supplementary Figure S12.**
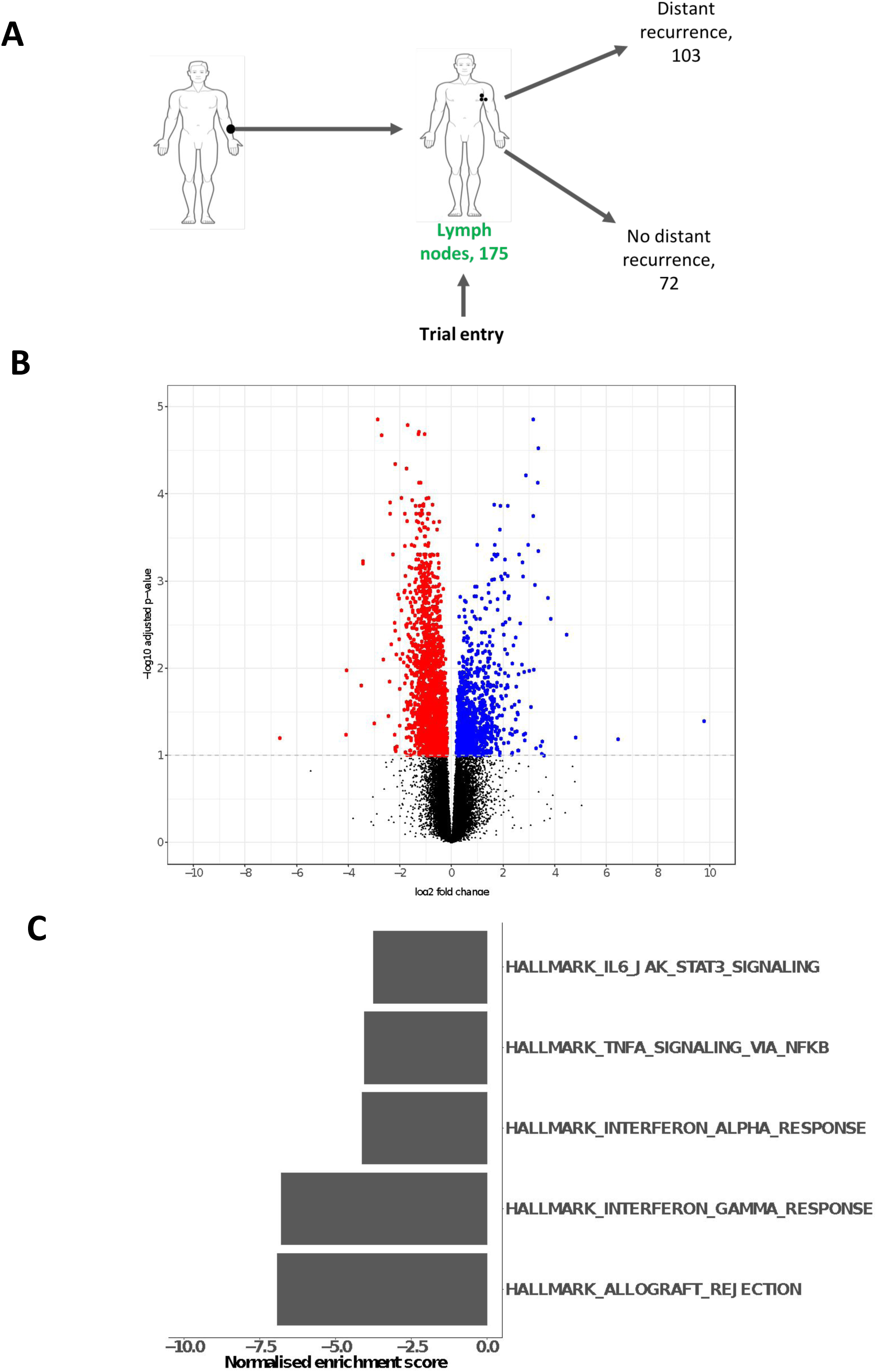
Results of differential expression and gene-set enrichment analysis comparing metastases vs non-metastases in lymph node samples, uncovering the same results as those obtained from skin samples. **A)** A schematic of the number of samples in the covariate corrected differential expression analysis. **B)** Volcano plot showing more downregulated (red points; 1967/3022 (65.1%)) than upregulated DEGs (blue points; 1055/3022 (34.9%)). **C)** Barplot showing downregulation of the same top five immune-related pathways as those identified in skin sample analyses (FDR corrected p-value<0.01 for all gene sets) (supplementary figure S11A).

**Supplementary Figure S13.**
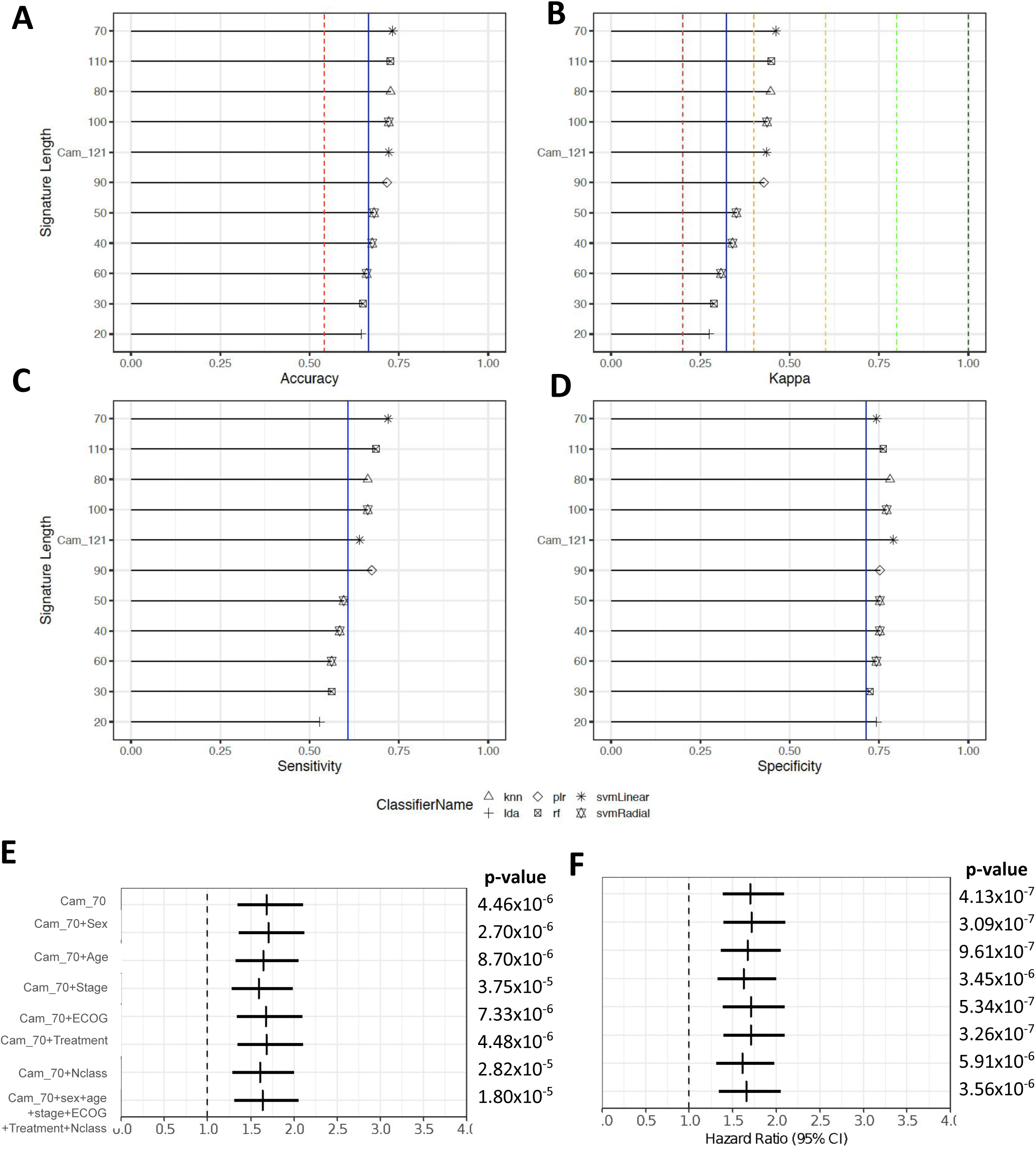
Feature reduction analysis showing signature length was reduced to 70 genes without compromising performance in our dataset but failed in external dataset. Barplots indicating, for each signature length (y-axes), the scores of **A)** accuracy, **B)** kappa, **C)** sensitivity and **D)** specificity (x-axes) of the top performing classifier (among the 8 considered). The blue line represents the corresponding scores of the top performing classifier (in terms of accuracy and kappa) on baseline clinical covariates. The dashed red line in (A) represents the accuracy achieved when the classifier only predicts the majority class (metastases=“No” in our case, 105/194 (54.12%)). Coloured lines in (B) represent the range of agreement bands of [cite whoever] defined for Cohen’s Kappa values. Forest plot indicating the hazard ratio estimates (vertical bars) and 95% confidence intervals (horizontal bars) related to the signature “Cam_70” when predicting **E)** OS and **F)** PFS by means of Cox proportional hazard models while controlling for different (sets of) clinical variables (y-axes). The Wald t-test p-values corresponding to the signature “Cam_70” parameter in each model are indicated on the right.

**Supplementary Figure S13.**
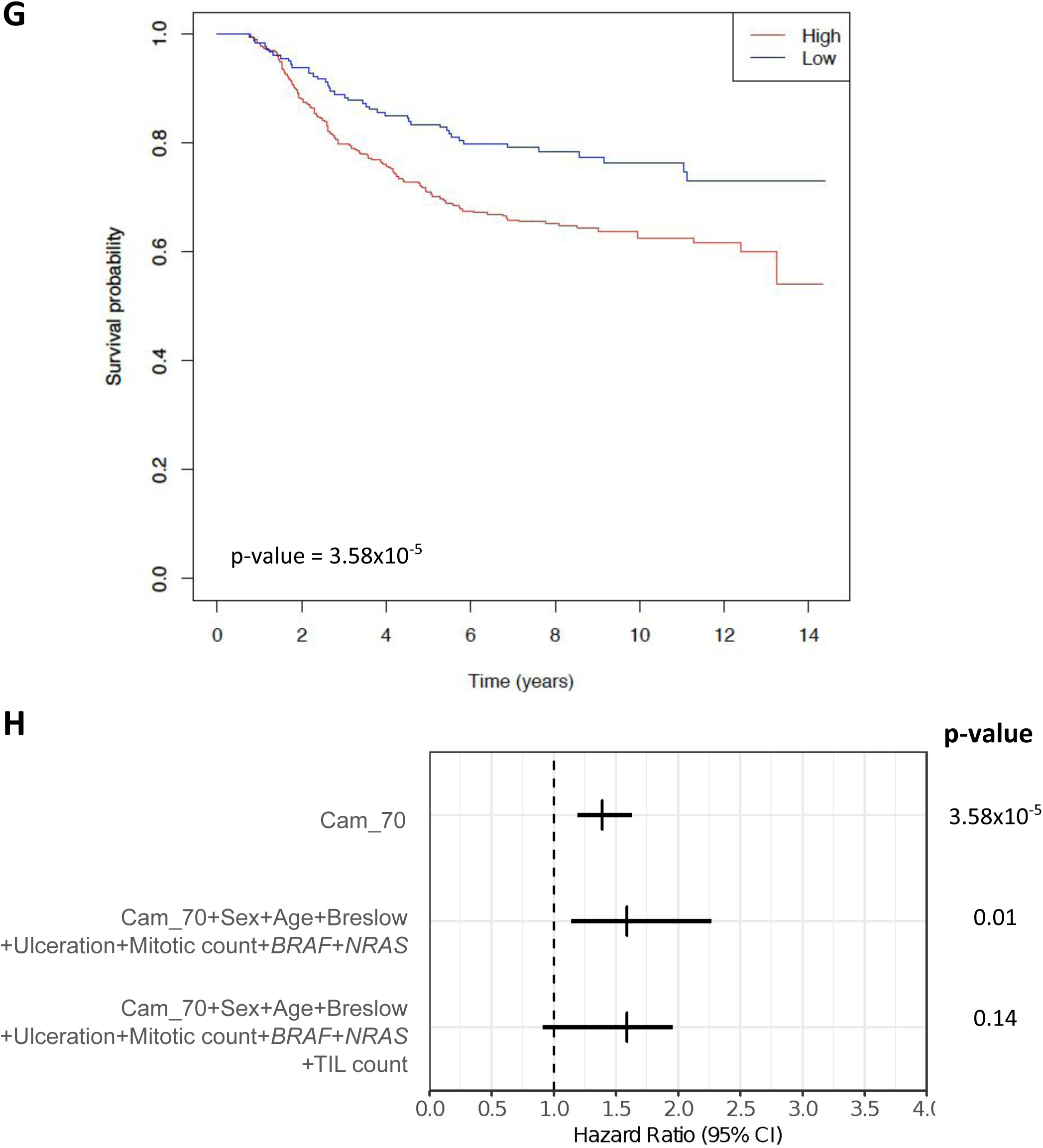
Feature reduction analysis showing signature length was reduced to 70 genes without compromising performance in our dataset but failed in external dataset. Validation on independently acquired external dataset (Leeds Melanoma Cohort, n = 677 samples): **G)** K-M curve comparing the Melanoma specific survival probabilities (y-axis) as a function of time (x-axis) of groups with high and low “Cam_70” (quantile 0.33 split) **H)** Forest plot indicating the hazard ratio estimates (vertical bars) and 95% confidence intervals (horizontal bars) related to the signature “Cam_70” when predicting Melanoma specific survival by means of Cox proportional hazard models while controlling for different (sets of) clinical variables (y-axes). Multivariate correction was undertaken for sex, age, breslow thickness, ulceration, mitotic count as well as *BRAF* and *NRAS* mutation status. In the final row, correction was also undertaken for the tumour infiltrating lymphocyte (TIL) score which diminished the prognostic value of the signature. The Wald t-test p-values corresponding to the signature “Cam_70” parameter in each model are indicated on the right.

**Supplementary Figure S14.**
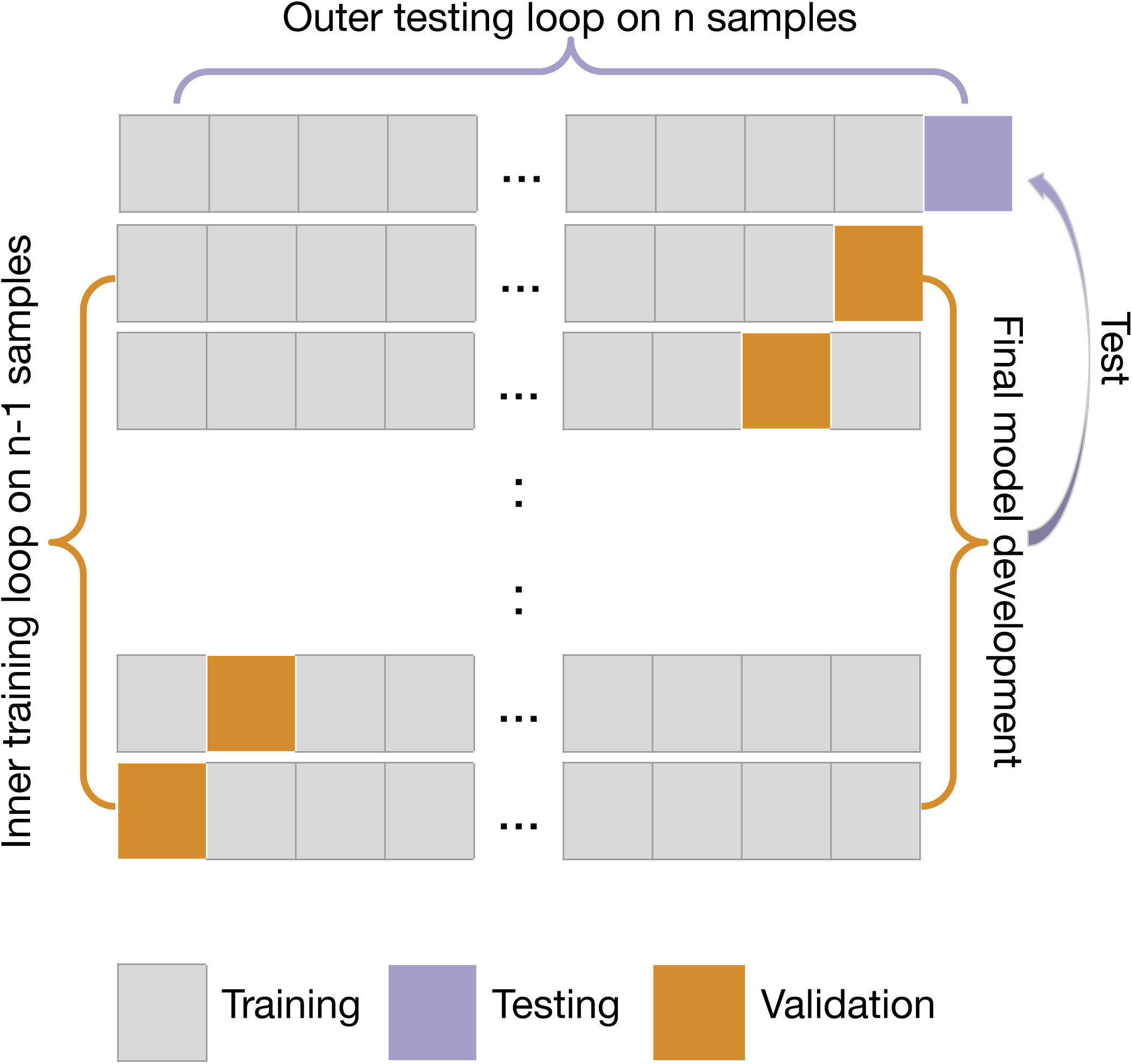
Schematic description of nested leave-one-out cross-validation (LOOCV). Outer test loop (Row 1): Each observation is iteratively considered as a test-set (violet observation) while the 193 remaining ones constitute the training set (grey observations). Nested inner loop in the training set (Rows 2 to last): Each observation of the training set is then iteratively considered as a validation set (orange observation), aiming to assess the properties (accuracy, kappa) of models and classifiers trained on the remaining 192 observations. At this level, 100 random combinations of hyperparameters are used for each classifier. The model and set of leading to the highest average criterion (accuracy, kappa) on the validation is then used on the outer loop test-set.

## Notes

**Author conflicts of interests:** PC is an advisory board member at Roche. MT/YY are employees and stockholders at Roche/Genentech. DJA is a consultant at Microbiotica

## References

1. Siegel RL, Miller KD, Jemal A. Cancer statistics, 2018. CA Cancer J Clin 2018;68(1):7–30 doi 10.3322/caac.21442.

2. Spain L, Larkin J, Turajlic S. New survival standards for advanced melanoma. Br J Cancer 2020 doi10.1038/s41416-020-0738-5.

3. Whiteman DC, Green AC, Olsen CM. The Growing Burden of Invasive Melanoma: Projections of Incidence Rates and Numbers of New Cases in Six Susceptible Populations through 2031. J Invest Dermatol 2016;136(6):1161–71 doi 10.1016/j.jid.2016.01.035.

4. Tarhini AA, Lee SJ, Hodi FS, Rao UNM, Cohen GI, Hamid O, et al. Phase III Study of Adjuvant Ipilimumab (3 or 10 mg/kg) Versus High-Dose Interferon Alfa-2b for Resected High-Risk Melanoma: North American Intergroup E1609. J Clin Oncol 2020;38(6):567–75 doi10.1200/jco.19.01381.

5. Robert C, Ribas A, Schachter J, Arance A, Grob JJ, Mortier L, et al. Pembrolizumab versus ipilimumab in advanced melanoma (KEYNOTE-006): post-hoc 5-year results from an open-label, multicentre, randomised, controlled, phase 3 study. Lancet Oncol 2019;20(9):1239–51 doi10.1016/s1470-2045(19)30388-2.

6. Maio M, Lewis K, Demidov L, Mandala M, Bondarenko I, Ascierto PA, et al. Adjuvant vemurafenib in resected, BRAF(V600) mutation-positive melanoma (BRIM8): a randomised, double-blind, placebo-controlled, multicentre, phase 3 trial. Lancet Oncol 2018;19(4):510–20 doi10.1016/s1470-2045(18)30106-2.

7. Long GV, Hauschild A, Santinami M, Atkinson V, Mandala M, Chiarion-Sileni V, et al. Adjuvant Dabrafenib plus Trametinib in Stage III BRAF-Mutated Melanoma. The New England journal of medicine 2017;377(19):1813–23 doi10.1056/NEJMoa1708539.

8. Luke JJ, Ascierto PA, Carlino MS, Gershenwald JE, Grob JJ, Hauschild A, et al. KEYNOTE-716: Phase III study of adjuvant pembrolizumab versus placebo in resected high-risk stage II melanoma. Future Oncol 2020;16(3):4429–38 doi 10.2217/fon-2019-0666.

9. Poklepovic AS, Luke JJ. Considering adjuvant therapy for stage II melanoma. Cancer 2019 doi10.1002/cncr.32585.

10. 1. 10. Bhutiani N, Egger ME, McMasters KM. Optimizing Follow-up Assessment of Patients with Cutaneous Melanoma. Ann Surg Oncol 2017;24(4):861–3 doi10.1245/s10434-017-5771-0.

11. Jonsson G, Busch C, Knappskog S, Geisler J, Miletic H, Ringner M, et al. Gene expression profiling-based identification of molecular subtypes in stage IV melanomas with different clinical outcome. Clin Cancer Res 2010;16(13):3356–67 doi10.1158/1078-0432.Ccr-09-2509.

12. Gerami P, Cook RW, Russell MC, Wilkinson J, Amaria RN, Gonzalez R, et al. Gene expression profiling for molecular staging of cutaneous melanoma in patients undergoing sentinel lymph node biopsy. J Am Acad Dermatol 2015;72(5):780–5.e3 doi10.1016/j.jaad.2015.01.009.

13. Zager JS, Gastman BR, Leachman S, Gonzalez RC, Fleming MD, Ferris LK, et al. Performance of a prognostic 31-gene expression profile in an independent cohort of 523 cutaneous melanoma patients. BMC Cancer 2018;18(1):130 doi10.1186/s12885-018-4016-3.

14. Gastman BR, Gerami P, Kurley SJ, Cook RW, Leachman S, Vetto JT. Identification of patients at risk for metastasis using a prognostic 31-gene expression profile in subpopulations of melanoma patients with favorable outcomes by standard criteria. J Am Acad Dermatol 2018 doi https://doi.org/10.1016/j.jaad.2018.07.028.

15. Vetto JT, Hsueh EC, Gastman BR, Dillon LD, Monzon FA, Cook RW, et al. Guidance of sentinel lymph node biopsy decisions in patients with T1-T2 melanoma using gene expression profiling. Future Oncol 2019;15(11):1207–17 doi10.2217/fon-2018-0912.

16. Thakur R, Laye JP, Lauss M, Diaz JMS, O’Shea SJ, Poźniak J, et al. Transcriptomic analysis reveals prognostic molecular signatures of stage I melanoma. Clin Cancer Res 2019:clincanres.3659.2018 doi10.1158/1078-0432.Ccr-18-3659.

17. Swetter SM, Tsao H, Bichakjian CK, Curiel-Lewandrowski C, Elder DE, Gershenwald JE, et al. Guidelines of care for the management of primary cutaneous melanoma. J Am Acad Dermatol 2019;80(1):208–50 doi10.1016/j.jaad.2018.08.055.

18. Corrie PG, Marshall A, Nathan PD, Lorigan P, Gore M, Tahir S, et al. Adjuvant bevacizumab for melanoma patients at high risk of recurrence: survival analysis of the AVAST-M trial. Ann Oncol 2018;29(8):1843–52 doi10.1093/annonc/mdy229.

19. Balch CM, Gershenwald JE, Soong SJ, Thompson JF, Atkins MB, Byrd DR, et al. Final version of 2009 AJCC melanoma staging and classification. J Clin Oncol 2009;27(36):6199–206 doi10.1200/jco.2009.23.4799.

20. Harbst K, Staaf J, Lauss M, Karlsson A, Masback A, Johansson I, et al. Molecular profiling reveals low- and high-grade forms of primary melanoma. Clin Cancer Res 2012;18(15):4026–36 doi10.1158/1078-0432.Ccr-12-0343.

21. Venet D, Dumont JE, Detours V. Most random gene expression signatures are significantly associated with breast cancer outcome. PLoS Comput Biol 2011;7(10):e1002240.doi10.1371/journal.pcbi.1002240.

22. Angelova M, Charoentong P, Hackl H, Fischer ML, Snajder R, Krogsdam AM, et al. Characterization of the immunophenotypes and antigenomes of colorectal cancers reveals distinct tumor escape mechanisms and novel targets for immunotherapy. Genome Biol 2015;16(1):64 doi10.1186/s13059-015-0620-6.

23. van Houdt IS, Sluijter BJ, Moesbergen LM, Vos WM, de Gruijl TD, Molenkamp BG, et al. Favorable outcome in clinically stage II melanoma patients is associated with the presence of activated tumor infiltrating T-lymphocytes and preserved MHC class I antigen expression. Int J Cancer 2008;123(3):609–15 doi10.1002/ijc.23543.

24. Krynitz B, Rozell BL, Lyth J, Smedby KE, Lindelof B. Cutaneous malignant melanoma in the Swedish organ transplantation cohort: A study of clinicopathological characteristics and mortality. J Am Acad Dermatol 2015;73(1):106–13.e2 doi10.1016/j.jaad.2015.03.045.

25. Thomas NE, Busam KJ, From L, Kricker A, Armstrong BK, Anton-Culver H, et al. Tumor-infiltrating lymphocyte grade in primary melanomas is independently associated with melanoma-specific survival in the population-based genes, environment and melanoma study. J Clin Oncol 2013;31(33):4252–9 doi10.1200/jco.2013.51.3002.

26. Azimi F, Scolyer RA, Rumcheva P, Moncrieff M, Murali R, McCarthy SW, et al. Tumor-infiltrating lymphocyte grade is an independent predictor of sentinel lymph node status and survival in patients with cutaneous melanoma. J Clin Oncol 2012;30(21):2678–83 doi10.1200/jco.2011.37.8539.

27. Donizy P, Kaczorowski M, Halon A, Leskiewicz M, Kozyra C, Matkowski R. Paucity of tumor-infiltrating lymphocytes is an unfavorable prognosticator and predicts lymph node metastases in cutaneous melanoma patients. Anticancer Res 2015;35(1):351–8.

28. Nsengimana J, Laye J, Filia A, O’Shea S, Muralidhar S, Pozniak J, et al. beta-Catenin-mediated immune evasion pathway frequently operates in primary cutaneous melanomas. J Clin Invest 2018;128(5):2048–63 doi10.1172/jci95351.

29. Pruessmann W, Rytlewski J, Wilmott J, Mihm MC, Attrill GH, Dyring-Andersen B, et al. Molecular analysis of primary melanoma T cells identifies patients at risk for metastatic recurrence. Nature Cancer 2020 doi10.1038/s43018-019-0019-5.

30. Vilain RE, Menzies AM, Wilmott JS, Kakavand H, Madore J, Guminski A, et al. Dynamic Changes in PD-L1 Expression and Immune Infiltrates Early During Treatment Predict Response to PD-1 Blockade in Melanoma. Clin Cancer Res 2017;23(17):5024–33 doi10.1158/1078-0432.Ccr-16-0698.

31. https://github.com/nunofonseca/fastq_utils [accessed August 2018].

32. http://www.bioinformatics.babraham.ac.uk/projects/fastqc/ [accessed August 2018].

33. Kim D, Pertea G, Trapnell C, Pimentel H, Kelley R, Salzberg SL. TopHat2: accurate alignment of transcriptomes in the presence of insertions, deletions and gene fusions. Genome Biol 2013;14(4):R36 doi10.1186/gb-2013-14-4-r36.

34. Anders S, Pyl PT, Huber W. HTSeq--a Python framework to work with high-throughput sequencing data. Bioinformatics 2015;31(2):166–9 doi10.1093/bioinformatics/btu638.

35. Hothorn T, Hornik K, van de Wiel MA, Zeileis. A Lego system for conditional inference. The American Statistician 2006;60(3):257–63.

36. Love MI, Huber W, Anders S. Moderated estimation of fold change and dispersion for RNA-seq data with DESeq2. Genome Biol 2014;15(12):550 doi10.1186/s13059-014-0550-8.

37. Therneau TM. 2015 A Package for Survival Analysis in R. <https://CRAN.R-project.org/package=survival>.

38. Zhu A, Ibrahim JG, Love MI. Heavy-tailed prior distributions for sequence count data: removing the noise and preserving large differences. Bioinformatics 2019;35(12):2084–92 doi10.1093/bioinformatics/bty895.

39. Max Kuhn. Contributions from Jed Wing SW, Andre Williams, Chris Keefer, Allan Engelhardt, Tony Cooper, Zachary Mayer, Brenton Kenkel, the R Core Team, Michael Benesty, Reynald Lescarbeau, Andrew Ziem, Luca Scrucca, Yuan Tang, Can Candan and Tyler Hunt. 2019 caret: Classification and Regression Training. <https://CRAN.R-project.org/package=caret>.

40. Gelman A, Jakulin A, Pittau MG, Su YS. A Weakly Informative Default Prior Distribution For Logistic And Other Regression Models. The Annals of Applied Statistics 2008;2(4):1360–83.

41. Friedman J, Hastie T, Tibshirani R. Regularization Paths for Generalized Linear Models via Coordinate Descent. J Stat Softw 2010;33(1):1–22.

42. Thomas C, Peter H. Nearest neighbor pattern classification. IEEE transactions on information theory 1967;13(1):21–7.

43. Fisher RA. The use of multiple measurements in taxonomic problems. Annals of eugenics 1936;7(2):179–88.

44. Park MY, Hastie T. Penalized Logistic Regression for Detecting Gene Interactions. Biostatistics 2008;9(1):30–50.

45. Breiman L. Random forests. Machine learning 2001;45(1):5–32.

46. Boser BE, Guyon IM, Vapnik VN. A training algorithm for optimal margin classifiers. 1992.

47. Chen H. 2018 VennDiagram: Generate High-Resolution Venn and Euler Plots. R package version 1.6.20. <https://CRAN.R-project.org/package=VennDiagram>.

48. Team RC. 2019 R: A Language and Environment for Statistical Computing.<https://www.R-project.org/>.

49. Maechler M, Rousseeuw P, Croux C, Todorov V, Ruckstuhl MA, Salibian-Barrera M, et al. 2019 robustbase: Basic Robust Statistics. <http://CRAN.R-project.org/package=robustbase>.

50. Subramanian A, Tamayo P, Mootha VK, Mukherjee S, Ebert BL, Gillette MA, et al. Gene set enrichment analysis: a knowledge-based approach for interpreting genome-wide expression profiles. Proc Natl Acad Sci U S A 2005;102(43):15545–50 doi10.1073/pnas.0506580102.

51. Mootha VK, Lindgren CM, Eriksson KF, Subramanian A, Sihag S, Lehar J, et al. PGC-1alpha-responsive genes involved in oxidative phosphorylation are coordinately downregulated in human diabetes. Nat Genet 2003;34(3):267–73 doi10.1038/ng1180.

52. Liberzon A, Birger C, Thorvaldsdottir H, Ghandi M, Mesirov JP, Tamayo P. The Molecular Signatures Database (MSigDB) hallmark gene set collection. Cell systems 2015;1(6):417–25 doi10.1016/j.cels.2015.12.004

