## Supplementary Methods for "Tumour gene expression signature in primary melanoma predicts long-term outcomes: A prospective multicentre study"

#### 1. **Visualization of inherent distribution of samples**

To visualize if the samples cluster by their metastatic status, principal component analysis (PCA) was performed on the primary melanoma samples (n=194) and lymph node samples (n=177) using 1000 most variable genes. This analysis was performed using the prcomp function, plotted using the qplot function from ggplot2 package (1)(v 3.2.1) and the scree plot was generated using the *screeplot* function from R-package stats (2)(v 3.6.2). The samples were further colored by whether they metastasized or not and shaped by their tissue of origin.

#### 2. **Testing signature performance against randomly selected genes**

To make sure that our signature does better than randomly-selected genes both in terms of metastases prediction and OS/PFS survival, random genes of the same length as the signature of interest were selected from 19434 protein-coding genes and the analysis was repeated using the exact same pipeline to compare its performance against our signature. This process was repeated 1000 times without replacement using 1000 different seeds and the p-value suggesting the significance of our signature was calculated as a left-tailed event for predicting OS/PFS survival (in terms of p-value). This analysis was inspired from the SigCheck package (3)(v2.14.0) in R (v3.5.1) and it's plotting function sigCheckPlotSurvival was modified to accept the scores generated from our analysis.

#### 3. **Feature reduction analysis**

Reduction analysis was performed to assess whether our Cam_121 gene signature could be reduced to a smaller list of genes, without compromising prognostic accuracy. The genes in the Cam_121 gene signature were sorted in decreasing effect size (determined by the shrinked log fold change values) as shown in Supplementary table S2 and in each iteration 10 genes with the lowest effect size were removed from the analysis. Multiple classifiers (n=8) were trained using the expression data of the remaining genes in the signature and performance metrics were calculated for evaluation. The reduced signature giving the highest kappa values was considered for further analysis.

### **References**

1.	Wickham H. ggplot2: Elegant Graphics for Data Analysis. Springer-Verlag New York; 2016.

2.	Team RC. 2019 R: A Language and Environment for Statistical Computing. <<https://www.R-project.org/>.>.

3.	Stark R, Norden J. SigCheck: Check a gene signature's prognostic performance against random signatures, known signatures, and permuted data/metadata. **2019**.
